# Estimating Waiting Distances Between Genealogy Changes under a Multi-Species Extension of the Sequentially Markov Coalescent

**DOI:** 10.1101/2022.08.19.504573

**Authors:** Patrick F. McKenzie, Deren A. R. Eaton

## Abstract

Genomes are composed of a mosaic of segments inherited from different ancestors, each separated by past recombination events. Consequently, genealogical relationships among multiple genomes vary spatially across different genomic regions. Expectations for the amount of genealogical variation among unlinked (uncorrelated) genomic regions is well described for either a single population (coalescent) or multiple structured populations (multispecies coalescent). However, the expected similarity among genealogies at linked regions of a genome is less well characterized. Recently, an analytical solution was derived for the expected distribution of waiting distances between changes in genealogical trees spatially across a genome for a single population with constant effective population size. Here we describe a generalization of this result, in terms of the expected distribution of waiting distances between changes in genealogical trees and topologies, for multiple structured populations with branch-specific effective population sizes (i.e., under the multispecies coalescent). Our solutions establish an expectation for genetic linkage in multispecies datasets and provide a new likelihood framework for linking demographic models with local ancestry inference across genomes.

## 1 Introduction

The multispecies coalescent (MSC) is an extension of the coalescent (Kingman, 1982), a model that describes the distribution of genealogical histories among gene copies from a set of sampled individuals. Whereas the coalescent models a single panmictic population, the MSC includes constraints that prevent samples in different lineages from sharing a most recent genealogical ancestor until prior to a population divergence event that separates them (Maddison, 1997; Maddison & Knowles, 2006). Conceptually, the MSC can be viewed as a piecewise model composed of the standard coalescent applied to each interval of a “species tree” representing the relationships and divergence times among isolated lineages. Genealogies are constrained to be embedded within species trees (Fig. 1a), and the joint likelihood of MSC model parameters can be estimated from the coalescent times among a distribution of sampled genealogies (Degnan & Rosenberg, 2009; Rannala & Yang, 2003). In both the coalescent and MSC models, effective population size (*N*_*e*_) is the key parameter determining the rate of coalescence and can vary across different lineages.

**Figure 1.**
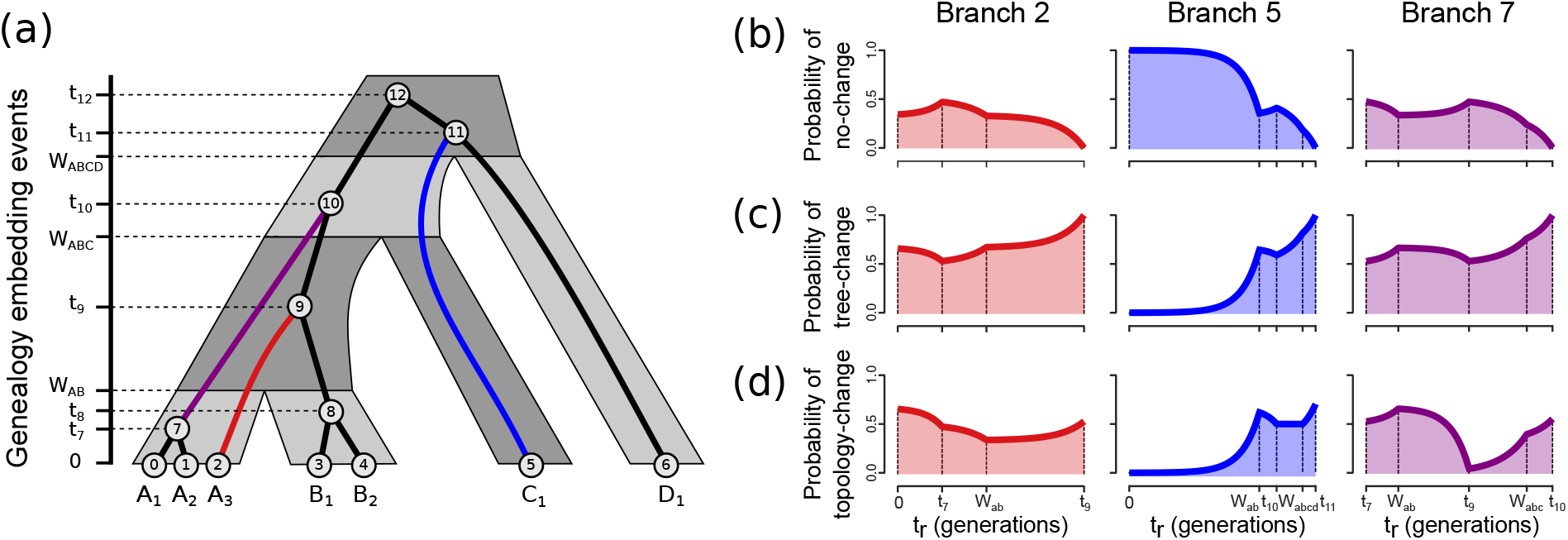
The MS-SMC models the probability of different recombination outcomes given a genealogy embedded in a multispecies coalescent (MSC) model. (a) A parameterized MSC model is composed of multiple discrete population intervals with associated effective population sizes (*N*_*e*_) and divergence times (*W*). Given an embedded genealogy, each interval can be further divided at coalescence events (e.g., *t*_9_) into a series of smaller intervals with constant coalescent rates. The parameters of these intervals make up a genealogy embedding table (see Table 1). (b-d) A recombination event leads to one of three possible outcomes between the genealogies in two sequential intervals of a genome: *no-change, tree-change*, or *topology-change*. The probabilities of each outcome as calculated under the MS-SMC are shown for three arbitrary branches. These probabilities are a function of the time (*t*_*r*_) and branch on which recombination occurs.

Importantly, both the coalescent and MSC are models of the expected distribution of *unlinked* (uncorrelated) genealogies. By contrast, two linked genealogies that are drawn from nearby regions of a genome are expected to be more similar than two random draws under these models. This spatial autocorrelation is a consequence of shared ancestry among samples at nearby regions, which decays over time and distance as recombination events reduce their shared ancestry. As a data structure, an ordered series of linked genealogies and the lengths of their associated intervals on a chromosome (Fig. 2) is represented by an ancestral recombination graph (ARG) (Griffiths & Marjoram, 1996), or similarly, a tree-sequence (Kelleher *et al*., 2016) (hereafter we will refer to it generically as an ARG).

**Table 1.**
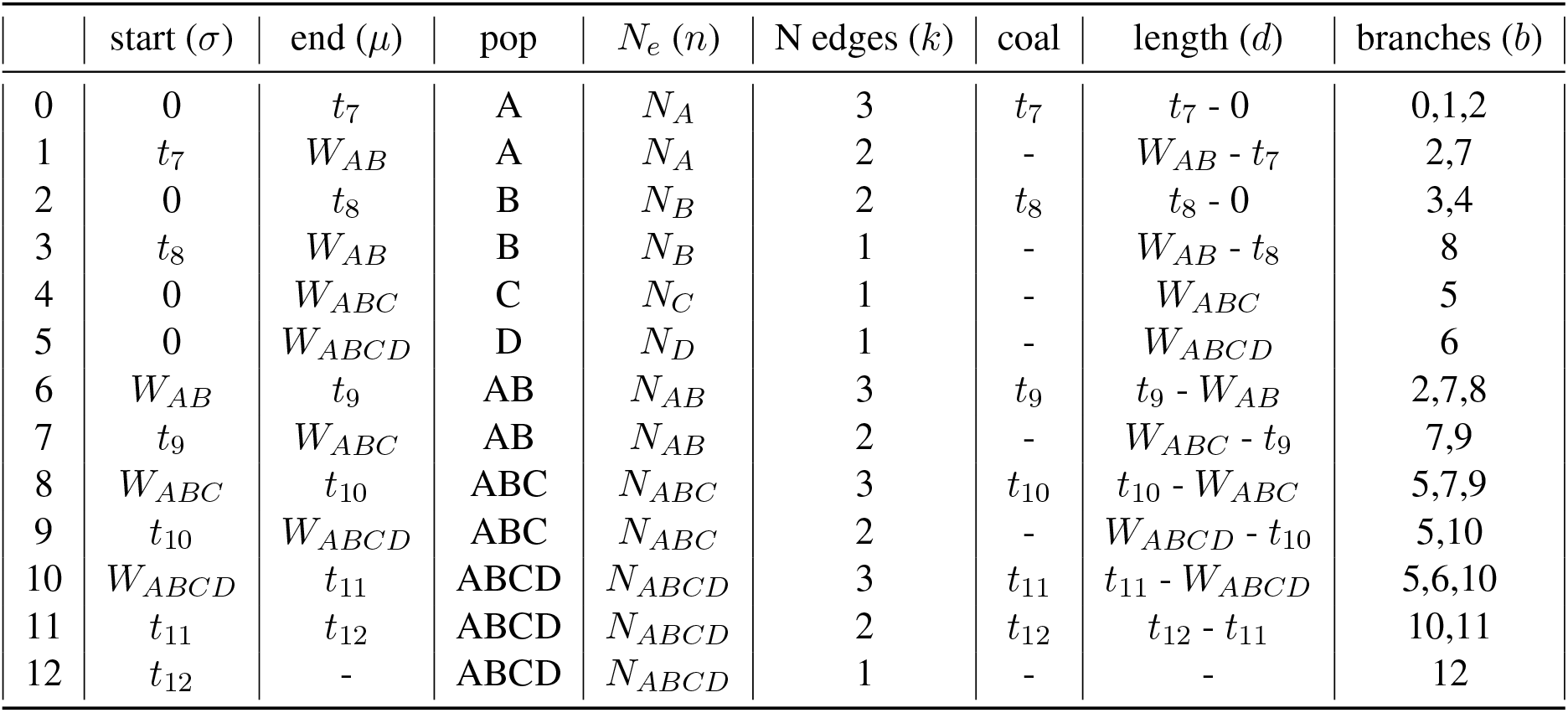
A genealogy embedding table for the genealogy and species tree in Figure 1. The probability of coalescence is constant within each discrete interval and is scaled by the number of lineages (*k*), the effective population size (*n*), and the interval length (*d*). Under the MS-SMC, the probability that a detached lineage will re-coalesce on a specific branch of the genealogy (e.g., branch 7 from Figure 1) is calculated using the piecewise constant probabilities from each discrete interval spanning that branch (e.g., rows 1, 6, 7, and 8).

**Figure 2.**
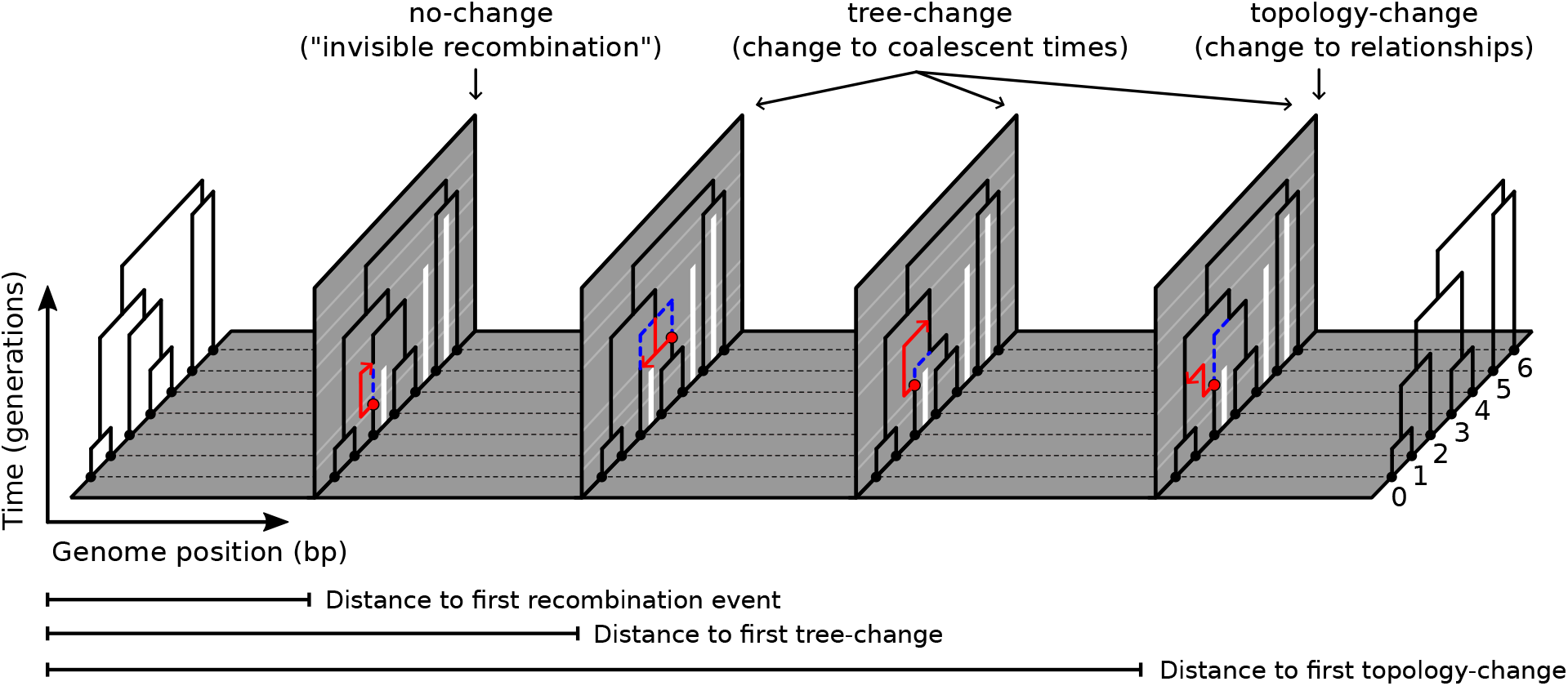
An ancestral recombination graph (ARG) is composed of a series of genealogies each spanning non-overlapping intervals of a genome separated by recombination breakpoints. Here, four recombination breakpoints separate the intervals’ start and end positions along a small chromosome alignment. The individual genealogies represent the history of a set of 7 samples constrained by a 4-tip species tree model (as in Fig. 1). Recombination events are indicated by vertical panels. Each shows the SMC’ process by which a subtree (below the red circle) is detached and then re-coalesces (red arrow) with one of the remaining lineages. The former edge (blue dotted) which existed throughout the interval to the left of a panel is replaced by a new edge (red) through the subsequent interval. Vertical bars (white) represent barriers to coalescence between samples in different species tree intervals (MSC model lineages). Four categories of recombination events are shown from left to right, representing different outcomes based on the lineage with which a detached subtree re-coalesces. These are grouped more generally into three *event types*: (1) no-change, (2) tree-change, and (3) topology-change. Every recombination event causes either a no-change or tree-change event, whereas topology-change events are a subset of possible tree-change events. The expected waiting distance until a specific recombination event type occurs can be calculated under the MS-SMC given a starting genealogy, MSC model, and recombination rate.

An algorithm to stochastically simulate sequences under the coalescent with recombination was developed early on (Hudson, 1983) and was later extended as a spatial algorithm for stochastically generating full ARGs (Wiuf & Hein, 1999). This process models the difference between sequential genealogies by randomly detaching an edge from a genealogy and sampling a waiting time (based on the population coalescent rate) until it reconnects to the genealogy at a different shared ancestor. Thus, under this spatial model, a set of samples effectively substitutes one ancestor for another on either side of each recombination breakpoint. Implicit to this algorithm is the assumption that recombination occurs at some predictable rate (or rate map) from which the expected waiting distance between recombination events can be modeled as an exponentially distributed random variable (Wiuf & Hein, 1999). Although generating ARGs consistent with a demographic model is relatively simple under this process, inferring an ARG from sequence data – composing a set of recombination breakpoints and local genealogy inferences – remains highly challenging (Brandt *et al*., 2022). This stems in part from the great complexity of this problem but also reflects limitations of our current models for extracting historical information from linked genome data.

A major advance was achieved through development of the sequentially Markov coalescent (SMC), a simpler approximation of the coalescent with recombination that restricts the types of recombination events that can occur (McVean & Cardin, 2005). Specifically, an edge that is detached from a genealogy by recombination is allowed only to re-coalesce with ancestral lineages that contributed genetic material to samples in that interval (as opposed to re-coalescing with any ancestral lineage). This greatly reduces the space of possible ARGs without changing the expected distribution of sequential genealogies, and in doing so it enables modeling changes between sequential genealogies as a Markov process (McVean & Cardin, 2005). Under these assumptions a tractable likelihood framework can be developed. Because neither genealogies nor segment lengths can be observed directly, most SMC-based inference methods use a hidden Markov model (HMM) to treat these as hidden states that influence observable changes in sequence data (Spence *et al*., 2018). Examples of inference tools built on the SMC framework include PSMC (Li & Durbin, 2011) and MSMC (Schiffels & Durbin, 2014) which use pairwise coalescent times between sequential genealogies to infer changes in effective population sizes through time, and ARG-weaver (Hubisz & Siepel, 2020; Rasmussen *et al*., 2014), which infers ARGs from genome alignments using an SMC-based conditional sampling method.

Marjoram & Wall (2006) described an important extension to the SMC, termed the SMC’, for additionally modeling “invisible” recombination events, in which a detached lineage re-attaches with its own ancestral lineage prior to the time of that lineage’s next coalescent event. This leads to no change between the genealogies in two sequential genomic intervals despite the occurrence of a recombination event between them. The inclusion of such events has been shown to significantly improve inference methods (Wilton *et al*., 2015). Under the SMC’, a detached lineage can thus re-coalesce with an allowable ancestral lineage over a continuous range of time, leading to one of four possible categorical patterns for the relationship between two sequential genealogies (Fig. 2, Fig. S1): (1) no change (2) shortening of a coalescent time; (3) lengthening of a coalescent time; and (4) a change to the genealogical topology (relationships). These can be grouped more generally into three types: “no-change” (category 1), “tree-change” (categories 2-4), and “topology-change” (category 4). Recently, Deng *et al*. (2021) derived a set of solutions for the expected waiting distances to each of these three types of outcomes for a single population with constant effective population size. This provided an important advance by establishing a neutral expectation not only for the distance until the next recombination event occurs, but more specifically, for the distance until different categorical types of recombination events occur. Such events leave different detectable signatures in sequence data and extend across different spatial distances of the genome, thus extending the scale over which information from spatial genealogical patterns can be extracted from genomes.

Here, we extend the methods of Deng *et al*. (2021) to an MSC framework to estimate the expected waiting distances between different types of genealogy changes under a parameterized species tree model. The waiting distances between recombination events that cause topology-change may be of greatest interest, as they leave the most detectable signatures in sequence data and are relevant to expected gene tree distributions that form an important component of many MSC-based methods (Baum, 2007; Degnan & Rosenberg, 2009; Knowles & Kubatko, 2011). The waiting distance between topology-change events may include multiple intervening recombination events of the no-change or tree-change type (Fig. 2). The relative occurrence of events that do not result in topology changes can be especially high in small sample sizes (Wilton *et al*., 2015), which are common in MSC-type datasets in which samples are partitioned among species tree intervals. Since the partitioning of coalescent events among species tree intervals is expected to constrain the types of recombination events that will be observed, the distributions of waiting distances between different types of genealogy changes should be highly dependent on, and thus informative about, the species tree model. We refer to the general framework of embedding the SMC’ in an MSC model as the MS-SMC.

## 2 Approach

### 2.1 Comparison to Deng *et al*. (2021)

Our approach is a generalization of the Deng *et al*. (2021) derivation of waiting distances to genealogy changes for a single population of constant size. We modified the single-population model to (1) include barriers to coalescence imposed by a species tree topology, and (2) integrate over changing coalescence rates along paths through multiple species tree intervals with different effective population sizes. In addition, we have implemented a novel likelihood framework for inferring parameters of an MSC model from observed waiting distances between genealogies.

### 2.2 MSC model description

Given an MSC model (*𝒮*) composed of a species tree topology with divergence times (*W*) in units of generations and constant diploid effective population sizes (*N*_*e*_) assigned to each branch, an embedded genealogy (*𝒢*) for any number of sampled gene copies (*k*) can be generated by randomly sampling coalescent times at which to join two samples into a common ancestor, starting from samples at the present in each interval. Following Kingman (1982), the probability of a coalescent event one generation in the past (reducing the number of samples from *k* to *k*-1) is given by equation 1. From this, we can model the expected waiting time (𝔼[*t*_*k*_]) until the next coalescence event as an exponentially distributed random variable with rate parameter *λ*_*k*_:

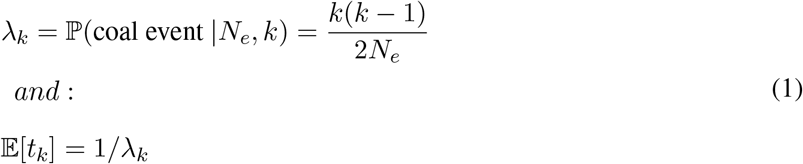

In a single population model with constant *N*_*e*_ the expected waiting time between coalescent events increases monotonically after each coalescent event, since the number of remaining samples always decreases. In an MSC model, however, the expected waiting time between coalescence events can increase or decrease through time, as the transition from one population interval to another can be associated with a different *N*_*e*_ value and an increase in the number of samples.

Based on this generative framework for sampling genealogies, likelihood solutions have been developed to fit coalescent model parameters, such as *N*_*e*_ in single population models (Kingman, 1982), or multiple *N*_*e*_ and *W* parameters in MSC models (Rannala & Yang, 2003), based on inferred coalescence times. In the latter framework, each species tree branch interval is treated independently, such that the likelihood of a genealogy embedding is calculated from the joint probability of observing each distribution of coalescent waiting times within each species tree branch interval. A key feature of these equations is that when *k* lineages are present, we can use the coalescent rate parameters (*λ*_*k*_) to calculate the likelihood of observed waiting times between coalescent events (*t*_*k*_) in each population interval from an exponential probability density function:

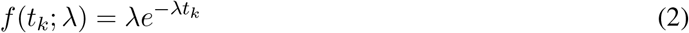

### 2.3 MS-SMC model description and notation

Under the SMC’ model, sampling of a linked genealogy requires considering not only demographic model parameters, as we did above, but also an existing genealogy – it is a method for sampling the next genealogy conditional on the previously observed one. If we define the previous genealogy as *𝒢*, and the sum of its edge lengths as *L*(*𝒢*), then under the assumption of a constant recombination rate through time, a recombination break point can be uniformly sampled from *L*(*𝒢*) to occur with equal probability anywhere on *𝒢*. A recombination event creates a bisection on a branch, separating a subtree below the cut from the rest of the genealogy (Fig. 2, Fig. S1). The subtree must then re-coalesce with an edge on the genealogy from which it was detached at a time above the recombination event. The waiting time until this re-coalescence event occurs is sampled stochastically with an expectation determined by the number of samples and the coalescent rate, similar to equation 1.

In a single population model with constant *N*_*e*_, the expected waiting time until re-coalescence increases monotonically with each coalescence event backwards in time, since each event decreases *k*. Once again, the MSC model differs from this: coalescent events similarly decrease *k*, but the merging of species tree branches into ancestral intervals increases *k*, and *N*_*e*_ can also vary among species tree intervals. Thus, the probability that a detached subtree re-coalesces to the genealogy can vary through time along its path of possible reconnection points through different species tree intervals (Fig. 1b-d). To calculate these probabilities, a species tree and genealogy can be decomposed into a series of relevant intervals between events that change rates of coalescence, which we refer to as the genealogy embedding table (Table 1). From this table it is possible to calculate the probabilities of different recombination event type outcomes and, consequently, to model the expected waiting distances until specific recombination event types occur.

Each interval of the genealogy embedding table contains a constant number of genealogy branches (*k*_*i*_) and a constant effective population size (*n*_*i*_) such that the rate of coalescence is also constant. We define a number of additional variables related to this table. The length of each interval is *d*_*i*_, and its lower and upper bounds are *σ*_*i*_ and *μ*_*i*_, respectively. Similarly, the lower and upper bounds of a genealogy branch (*b*) are defined as 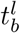 and 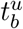, respectively. An indexing variable, *ℐ*_*b*_, is defined as the ordered set of intervals that are spanned by a specific branch of an embedded genealogy. As an example, consider branch 7 from Fig. 1. This branch spans four intervals, labeled as rows 1, 6, 7, and 8 in Table 1, and so *ℐ*_7_ = {1,6,7,8}. As we will demonstrate below, this and related indexing variables will be used to calculate the probabilities of re-coalescence in different intervals, and on different branches, based on their rates of coalescence. A summary of all variables in our notation is available in table S1.

### 2.4 Deriving probabilities of genealogy changes in the MS-SMC

A recombination event occurring on *𝒢* can result in three types of outcomes (Fig. 2). Of these, there is a zero-sum relationship between a no-change and tree-change event, such that one or the other must occur. Therefore, as a first step towards describing probability statements for each of these event types, we focus first on deriving the probability of a no-change event (also termed a tree-unchanged event; Fig. 3a), which is the simplest outcome. Then, from the law of total probability, we also have a result for the probability of recombination resulting in a tree-change event. Finally, to calculate the probability of a topology-change event, we first derive a statement for the probability of a topology-unchanged event (Fig. 3d), which is the union of a no-change event and a subset of tree-change events, where the detached lineage is restricted according to which ancestral lineages it can re-coalesce with. Full detailed derivations of all solutions below are available in the Appendix of the Supplementary Materials.

**Figure 3.**
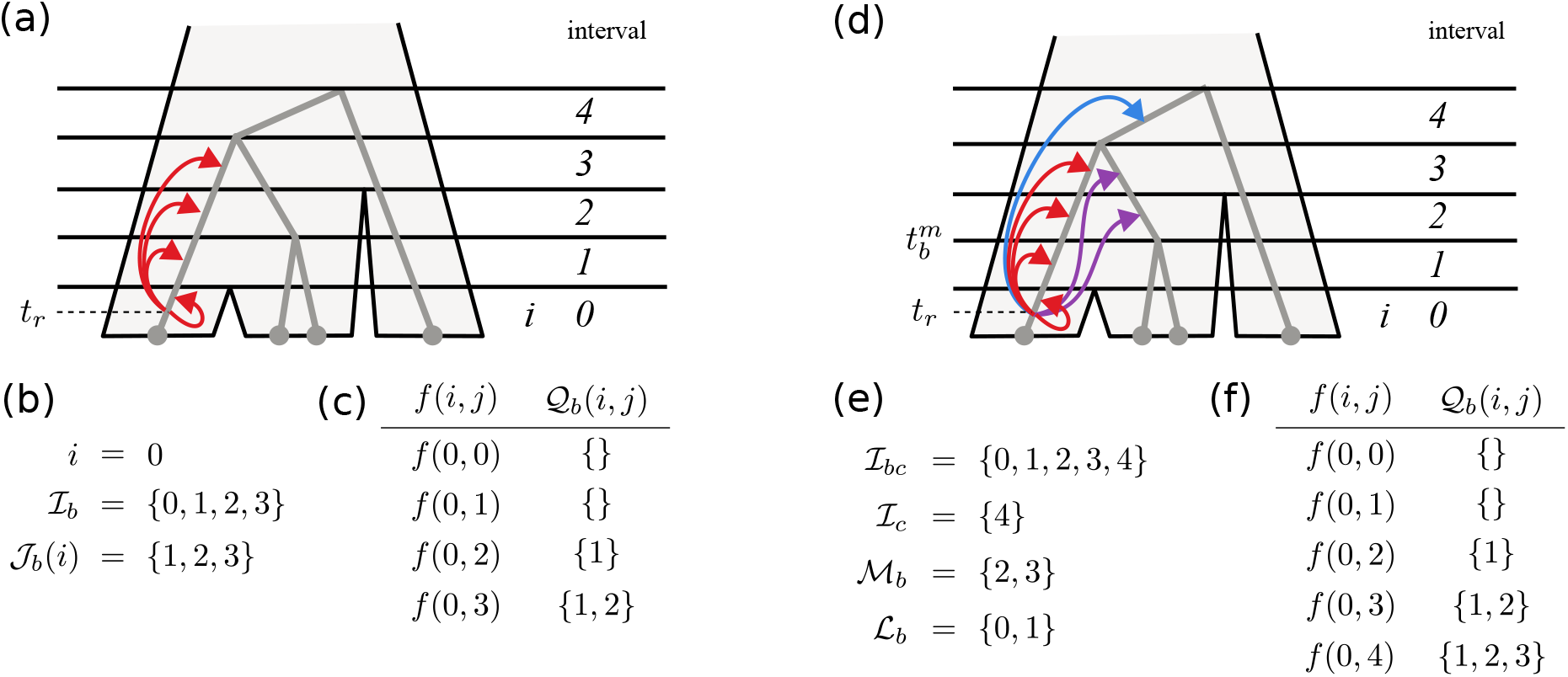
The probabilities of different recombination event types are calculated using piece-wise constant probabilities of recombination within discrete intervals. (a) The probability of a no-change event involves re-coalescing with the same branch on which recombination occurred, for which intervals are indexed using the variable *ℐ*_*b*_. (b) Indexing variables for calculating tree-change probabilities. The indexing variable *ℐ*_*b*_ records all intervals on branch *b*, and the recombination event for this example occurs in interval *i* = 0. *𝒥*_*b*_(*i*) returns all intervals in *ℐ*_*b*_ above interval *i*. (c,f) The function *f* (*i, j*) returns the probability that a subtree that detached in interval *i* will re-coalesce in interval *j*. This involves excluding the probability of re-coalescence in intervals between *i* and *j*, which are indexed as *𝒬*_*b*_(*i, j*). (d) The probability of a topology-unchanged event involves re-coalescing with the same branch on which recombination occurred, or with its parent or sibling branches. (e) Additional indexing variables return intervals on the parent branch (*ℐ*_*c*_), or branch *b* above (*ℳ*_*b*_) or below (*ℒ*_*b*_) the time (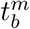) at which it overlaps in the same species tree interval as its sibling branch.

#### 2.4.1 Probability of a no-change event

We begin by assuming knowledge of when and where recombination takes place, in terms of a recombination event bisecting branch *b* at time *t*_*r*_. For no change to occur to the genealogy, the detached subtree must re-coalesce with its original branch – either in the same interval from which it detached, or in a later interval on the same branch (Fig. 3a). If it connects to any other lineage, this will cause a change to either the tree or topology. Equation 3 describes the probability of a no-change event given a genealogy embedded in a species tree and given the timing and branch on which recombination occurs. The interval in which recombination occurs is labeled *i*. The first two terms describe the probability that the subtree re-coalesces during interval *i* on branch *b* (i.e., *ii*), while the latter term is the probability of re-coalescing during a later interval on branch *b* (i.e., *ij*). For this, we define another indexing variable *𝒥*_*b*_(*i*) = *{j* ∈ *ℐ*_*b*_ | *j > i}*, for iterating over the ordered intervals above *i* on branch *b* (e.g., Fig. 3b).

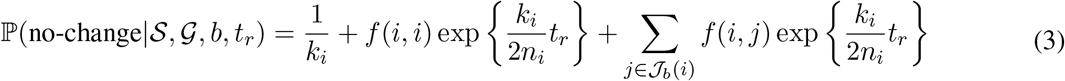

This equation is simplified by use of the function *f* (*i, j*) to return the piece-wise constant probabilities of re-coalescence between pairs of intervals. When *j* = *i*, this expression involves the probability of coalescing over the remaining length of interval *i* above *t*_*r*_; when *j > i* it involves the probability of coalescing in interval *j* and not coalescing in interval *i* or any other intervals between *i* and *j*. For this latter process, we define another indexing variable, *𝒬*_*b*_(*i, j*) = *{q* ∈ *ℐ*_*b*_ | *j > q > i}*, for iterating over the ordered intervals above *i* and below *j* on branch *b* (e.g., Fig. 3c). The function *f* (*i, j*) forms the core of the MS-SMC algorithm and will reappear in several later equations.

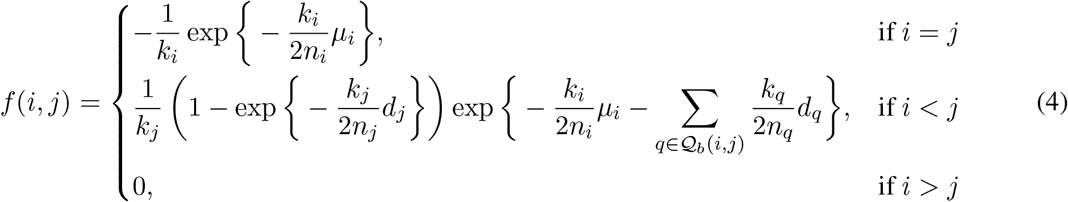

By integrating equation 3 across all times at which recombination could have occurred on branch *b* (assuming a uniform recombination rate through time) we obtain the probability that recombination anywhere on this branch does not change the tree:

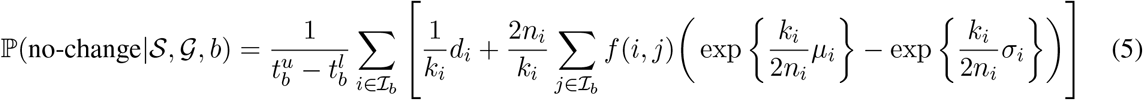

g

Finally, by summing across all branches on the tree while weighting each one by its relative proportion of edge length, we get the probability that a recombination event occurring anywhere on *𝒢* will result in a no-change event.

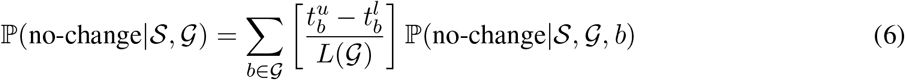

#### 2.4.2 Waiting distances to no-change and tree-change events

Under the SMC’, recombination is modeled as a Poisson point process such that the time between recombination events is exponentially distributed with rate parameter *λ*_*r*_: the product of the per-site per-generation recombination rate and summed branch lengths of the current genealogy (Wiuf & Hein, 1999) (equation 7). The likelihood of an observed distance (*x*) between recombination events spatially along the genome, in units of base pairs, can thus be calculated from the exponential probability density function (equation 8).

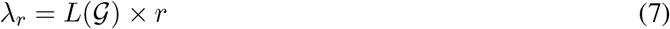

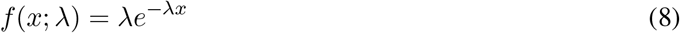

Having derived the probability that an individual recombination event is of the no-change type, we can now calculate the rate of no-change type events as a proportion of the rate of all types of recombination events. Here, waiting distances continue to be exponentially distributed. However, the new rate parameter *λ*_*n*_ is reduced proportionally by the probability that recombination causes no change to the genealogy (equation 9). Similarly, because a tree-change event is the opposite of a no-change event, its probability is one minus the probability of no-change (equation 10). This yields rate parameter *λ*_*g*_ for the exponential probability distribution of waiting distances between tree-change events.

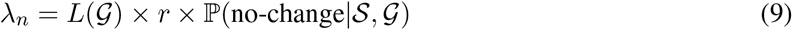

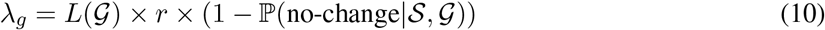

#### 2.4.3 Probability of topology-change

We next derive an analogous probability distribution for waiting distances between topology-change events. Similar to our approach for calculating tree-change probabilities as the opposite of those for a no-change (tree-unchanged) event, here we calculate topology-change probabilities as the opposite of those for a topology-unchanged event. Topology-unchanged events represent the union of all no-change events and the subset of possible tree-change events that only affect branch lengths but not the topology. Our approach for calculating these probabilities follows closely to that of Deng *et al*. (2021). In order to isolate re-coalescence events that do not change the topology, we must take into account which specific branches the detached subtree from branch *b* re-coalesces with. The relevant branches are its source (*b*), its sibling (*b*^*′*^), and its parent (*c*) (Fig. 3d). If the subtree re-coalesces with *b*, no change occurs; if it re-coalesces with *b*^*′*^, the topology remains the same but a coalescent time is shortened; and if it re-coalesces with *c*, the topology remains the same but a coalescent time is lengthened. A re-coalescence with any other branch will change the topology.

To index over relevant intervals across the three branches on which re-coalescence can occur, we define several additional variables. The lowest time point in which both *b* and *b*^*′*^ are present and exist within the same species tree interval is labeled 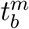. For a branch *b* with intervals *ℐ*_*b*_, the subset of intervals below 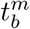 is *ℒ*_*b*_, and the subset above is *ℳ*_*b*_. The union of the sets of intervals on branches *b* and *c* is *ℐ*_*bc*_ (Fig. 3e).

Once again, we begin by assuming knowledge of the branch on which a recombination event occurs and of that event’s timing. This problem can be broken into two distinct cases: when *t*_*r*_ occurs below 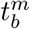, and when it occurs above 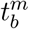. The former requires integrating over additional intervals on *b* that are not shared with *b*^*′*^. Over all intervals on the three relevant branches, these equations use the function *f* (*i, j*) to return the piecewise constant probabilities where recombination occurs in interval *i* and re-coalesces in interval *j* (Fig. 3f).

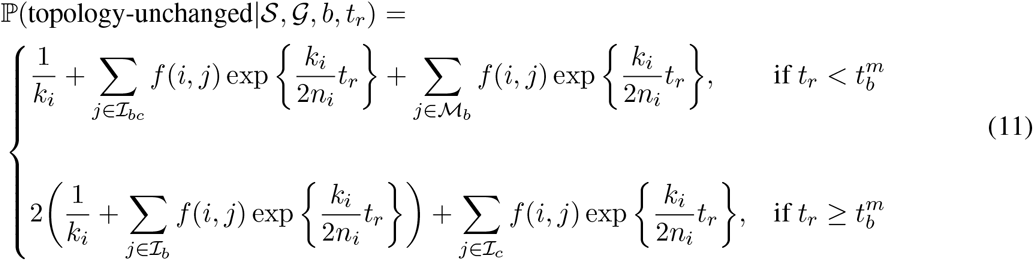

Next, the probability of a topology-unchanged event given recombination anywhere on a branch can be derived by integrating the previous equation over the entire length of a branch. Here, the terms *p*_*b*,1_ and *p*_*b*,2_ correspond to recombination occurring on branch *b* during a time that falls into either of the two cases in equation 11.

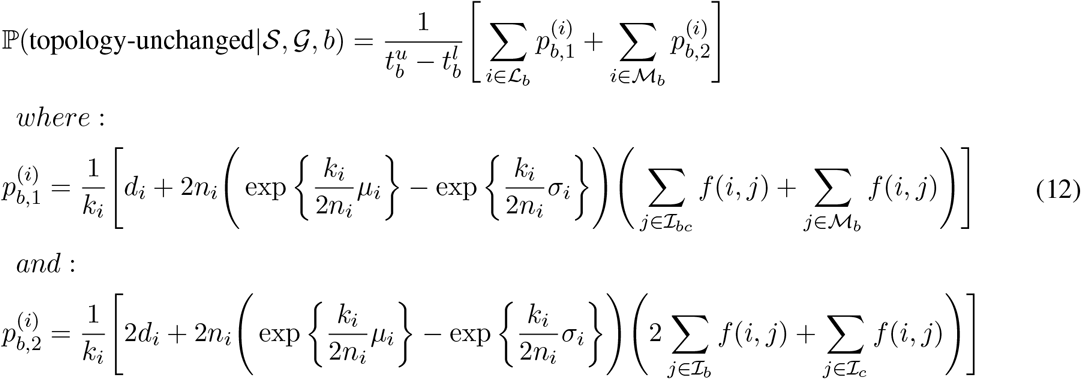

Finally, by summing equation 12 across all branches on a genealogy while weighting each by its proportion of summed branch lengths, we get the probability that a recombination event falling uniformly on the genealogy will result in a topology-unchanged event.

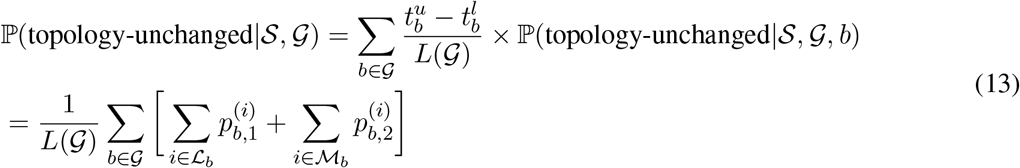

#### 2.4.4 Waiting distance to topology-change events

A recombination event either does or does not change the topology of a genealogy, and we can therefore get the probability of a topology-change event using our topology-unchanged probability statement. As with the previous waiting distance distributions, the distance between topology-change events given a parameterized MSC model can be modeled as an exponential probability distribution. Similar to how a rate parameter was derived for the distribution of waiting distances until a recombination event (equation 7), no-change event (equation 9), or tree-change event (equation 10), a rate parameter *λ*_*t*_ can be calculated from equation 13 for the probability of a topology-change event.

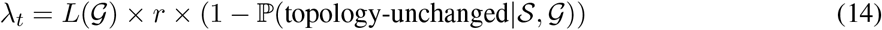

Unlike the exact solution for the expected waiting distance to a tree-change, the waiting distance for a topology-change is an approximation. This is because topology-change probabilities are not guaranteed to be homogeneous across some distance of the genome between topology-change events, since intermediate tree-change events could occur (e.g., the second and third recombination events in Fig. 2). We examine this and other potential sources of bias in our validations below and in the Supplementary Materials.

## 3 Results

### 3.1 Implementation

We have implemented our solutions for waiting distance calculations under the MS-SMC in the Python package *ipcoal* (McKenzie & Eaton, 2020a). This software includes functions that accept a parameterized MSC model and initial genealogy as input and return the probabilities of different recombination event types. These probabilities can be calculated for specific branches and times, for entire branches, or for entire genealogies. Functions are also available to calculate expected waiting distances and the likelihoods of observed waiting distances in an ARG using an exponential probability density parameterized by the MS-SMC. Our implementation is built upon *toytree* (Eaton, 2020), *scipy* (Virtanen *et al*., 2020), and *numpy* (Harris *et al*., 2020), and it uses jit-compilation with *numba* (Lam *et al*., 2015). Below we use *ipcoal* (v.0.4.0) to demonstrate the impact of MSC model parameters on waiting distances and to validate our solutions against expectations from coalescent simulations implemented in *msprime* (v.1.1.1) (Baumdicker *et al*., 2022). Source code is available at https://github.com/eaton-lab/ipcoal. Jupyter notebooks demonstrating the MS-SMC calculations and with reproducible code used for validations in this study are available at https://github.com/eaton-lab/waiting-distances.

### 3.2 Demonstration

Given a parameterized MSC model and initial genealogy, the probabilities of different types of recombination outcomes can be calculated and visualized as a function of when and where recombination occurs. This is demonstrated on an imbalanced 4-tip species tree with constant effective population size and with a genealogy of seven samples embedded, including three from lineage A, two from lineage B, and one from each of lineages C and D (Fig. 1a; Fig. S2). The probabilities of no-change, tree-change, or topology-change events, given a recombination event occurring on a branch at a particular time (equations 3 and 11, respectively) are shown for three selected branches on the example genealogy (Fig. 1b-d). Note that the probability of no-change and tree-change events are inversely related and sum to 1, since one or the other must occur at any recombination event. By contrast, a topology-change event is a subset of the probability of a tree-change; it is a tree-change event where the detached branch re-coalesces with a branch other than itself, its sibling, or its parent.

In general, the probability of a no-change event decreases, and the probability of a tree-change event increases, as recombination occurs closer to the top end of a branch (further back in time). This makes intuitive sense, since when recombination occurs at the top of a branch there is less time for it to re-coalesce with its same branch. Although this is a general trend, these probabilities do not behave monotonically along the length of a branch as they would in a single-population model with constant *N*_*e*_ (Deng *et al*., 2021). Instead, probabilities increase or decrease through the length of each interval as a function of the rates of coalescence in subsequent intervals and the probability that a detached lineage will re-coalesce in one of those intervals.

For example, consider branch 2, which exhibits an increase in the probability of no-change through its first branch interval from time 0 to *t*_7_ but then a decrease through the next interval from *t*_7_ to *W*_*AB*_ (Fig. 1b). The observed increase through the first interval is influenced by the fact that a re-coalescence in the subsequent interval is more likely to cause a no-change event, since that interval contains only two samples instead of three. By contrast, within the second interval, recombination events near the top are approaching the next species tree divergence event. After that event, the number of samples will increase back from 2 to 3, thus decreasing the probability of a no-change event. This visualization demonstrates how the probabilities of different recombination event types represent an integration over all the positions on a branch where recombination could occur, and all positions at or above each of these points (whether on the same or different available branches) where a detached subtree could re-coalesce.

Branch 7 provides a clear example for examining the probabilities of tree- and topology-change events. Of particular interest is the interval from *W*_*AB*_ to *W*_*ABC*_ where these probabilities diverge significantly (Fig. 1c-d). The probability of topology-change decreases faster than the probability of tree-change as recombination occurs closer to node *t*_9_. This is because following *t*_9_ there is a large stretch of time during which re-coalescence can only occur with the same branch or its sibling, neither of which can cause a topology-change event. It is only after *W*_*abc*_ that it is once again possible for re-coalescence to occur with a more distant branch that would result in topology-change. If the effective population size of this species tree interval (AB) were greater, then the probability of re-coalescence in a deeper interval would be more likely, and the probability of topology-change would decrease less severely near *t*_9_. This is true more generally, as can be seen by comparing edge probabilities across MSC models with different effective population sizes (Fig. S2). Effective population size affects the rate of re-coalescence and thus either smooths probabilities across intervals when *N*_*e*_ is high or accentuates differences among intervals when *N*_*e*_ is low.

### 3.3 Validation

To validate our analytical solutions for the probabilities of different recombination event outcomes and their associated waiting distances, we compared predictions of the MS-SMC with results from stochastic coalescent simulations. We set up three scenarios with increasing amounts of population structure: a single-population model, a two-population model, and an 8-tip phylogeny model (Fig. 4a). All analyses used a constant per-site per-generation recombination rate of 2e-9 and simulated tree sequences using the coalescent with recombination (i.e., the “hudson” ancestry model in msprime as opposed to the “smc_prime” model, which is an approximation), and the argument “record_full_arg=True” (to retain records of invisible recombination events). For each model we simulated genealogies for the same total number of samples (8, unless specified), divided evenly among lineages when models include multiple populations (Fig. 4a).

**Figure 4.**
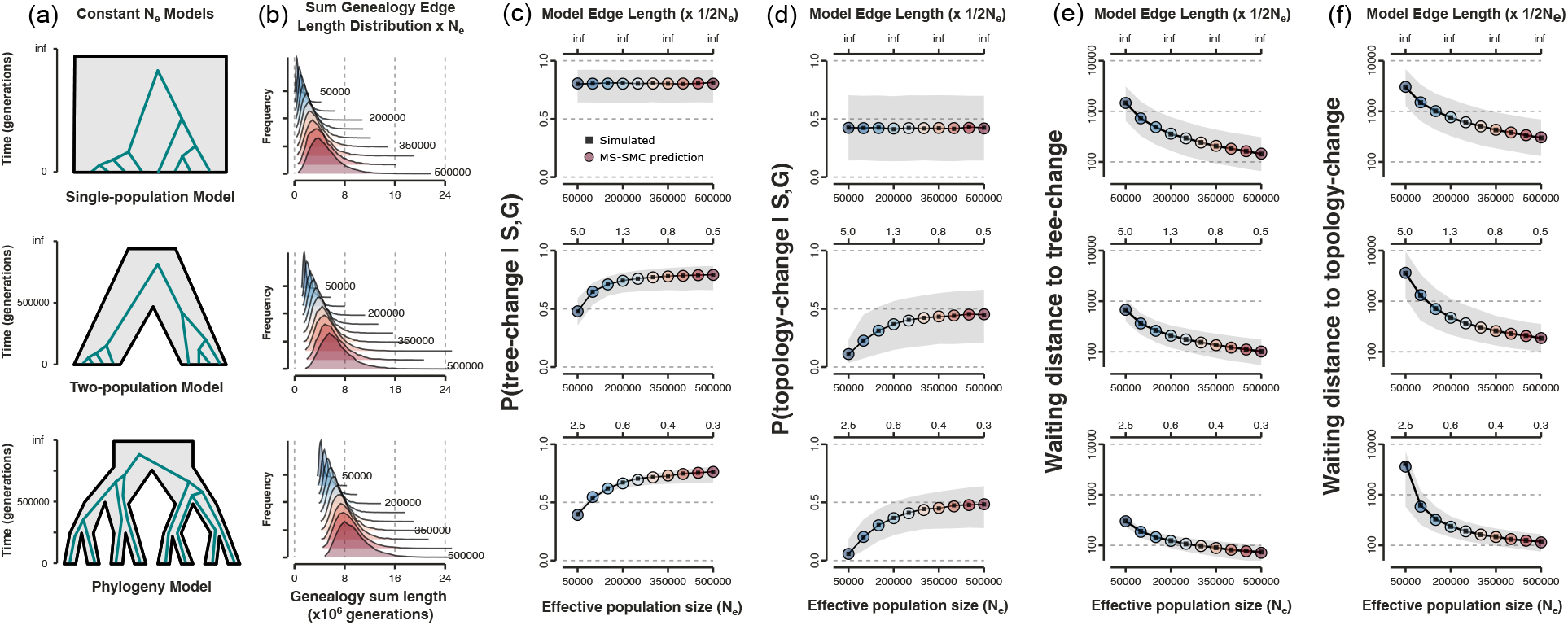
MS-SMC predictions validated against coalescent simulations. (a) Results are shown for three models containing 1, 2, or 8 populations. For each model, 10K tree sequences were simulated for 10 different constant *N*_*e*_ values between 50K and 500K. (b) The distributions of summed edge lengths of the first genealogy in each tree sequence. (c-d) The mean frequency (black square) with which the first observed recombination event was a tree-change (c) or topology-change (d) in a simulated tree sequence, and the mean (colored circle) and 95% CI (grey fill) of the predicted probability of tree or topology-change calculated from the first embedded genealogy in each tree sequence. Probabilities are constant with respect to *N*_*e*_ in the single population model but vary in models with population structure (also shown with respect to species tree interval lengths in coalescent units, across the top axis). (e-f) The mean waiting distance (black square) until the first observed tree-change (e) or topology-change (f) in a simulated tree sequence, and the mean (colored circle) and 95% CI (grey fill) of predicted waiting distances calculated using the first embedded genealogy in each tree sequence.

The exponential rate parameter (*λ*) for a probability distribution of waiting distances is a product of the per-site per-generation recombination rate (*r*), the sum of edge lengths on the current genealogy (*L*(*𝒢*)), and the probability (ℙ) of the specified event type (equations 9, 10, and 14). Across the three models examined, *r* remains constant, but both *L*(*𝒢*) and ℙ can vary due to population structure, where the effect of structure is scaled by *N*_*e*_. Therefore, we examined *L*(*𝒢*), ℙ, and the expected waiting distance calculated from their product, for each demographic model across a range of *N*_*e*_ values (Fig. 4b-f). For each value of *N*_*e*_, 10K tree sequences were simulated that included at least one topology-change event. The empirical probabilities of tree-change and topology-change events were then simply measured as the mean observed frequency at which the first recombination event in each tree sequence was a tree or topology change. Similarly, empirical waiting distances were measured as the mean distance until the first observation of each event type. These simulated expectations were then compared with analytical expectations computed under the MS-SMC, where probabilities and waiting distances were based only on the embedding of the first genealogy from each tree sequence into an MSC model parameterized to the specific constant *N*_*e*_ value.

Population structure enforces a lower limit on the length of coalescent times by requiring that genealogies can be embedded in a species tree. This has the effect of shifting both the minimum and mean of *L*(*𝒢*) higher (Fig. 4b). Because the per-generation recombination rate interacts with *L*(*𝒢*) (the opportunity over which recombination can occur) to determine the frequency of recombination, larger *L*(*𝒢*) induced by population structure will tend to decrease waiting distances between recombination events, all else being equal. However, all else does not remain equal. Population structure has a simultaneous opposing effect on waiting distances by decreasing the probability of tree or topology changes (ℙ; Fig. 4c-d), especially at low *N*_*e*_ values, where species tree constraints can make tree or topology changes unlikely to occur. This is a stark difference between the MS-SMC framework and a single population model: the probability of a tree- or topology-change event is strongly associated with *N*_*e*_ in the former but not affected at all in the latter. Consequently, the waiting distances between each event type in MSC models (Fig. 4e-f) represent a balance of the positive and negative effects of population structure on *L*(*𝒢*) and ℙ, respectively.

Our analytical predictions under the MS-SMC converge accurately on the mean results from stochastic coalescent simulations (Fig. 4c-f). Moreover, by examining the variance in these predictions with respect to MSC model parameters we further gain insights into the information contained in spatial genealogical patterns. For example, in the single population model there is high variance in both the probabilities of tree and topology changes, as well as in genealogy lengths, at any given *N*_*e*_ value. Consequently, waiting distances also exhibit high variance. Although waiting distances correlate with population *N*_*e*_ in this model, the differences in mean waiting distances are small relative to the variance. By contrast, multi-species models exhibit much less variance in predicted probabilities of tree or topology changes given a set of MSC model parameters (Fig. 4c-d) and also exhibit less variation in genealogy lengths. This leads to a stronger relationship between MSC model parameters and expected waiting distances (Fig. 4e-f), such that models with different parameters have less overlap in waiting distance expectations. Overall, this suggests that tree- and topology-change distances may contain more information in MSC models than in single population models.

### 3.4 Bias in waiting distance estimation

Estimated waiting distances under the MS-SMC harbor two potential sources of bias, the first stemming from assumptions of the SMC’ approximation, and the second from the approximate nature of waiting distance estimation for topology-change events. We examined both of these sources of error through comparison to stochastic simulations and found that their effects are generally small, and that multispecies models exhibit a similar magnitude of error as single population coalescent models (Supplementary Materials; Fig. S3, Fig. S4). The SMC’ approximation contributes the greatest error in all models, leading to a slight under-estimation of waiting distances. This is most observable when variance in waiting distances is high, as with topology-change waiting distances in models with low *N*_*e*_. Future work could seek to find a correction for this bias.

### 3.5 MS-SMC likelihood inference framework

The MS-SMC provides a statistical framework for predicting waiting distances until genealogy changes in an ARG given a parameterized MSC model. Because waiting distances between recombination events are exponentially distributed, we can use the exponential probability density (equation 8) as a likelihood function to describe the joint probability of observed waiting distances as a function of MSC model parameters. Applications of this approach could potentially provide improvements to ARG or species tree inference methods. Here, to explore the potential of this approach, we developed a maximum likelihood framework for the latter: we apply the MS-SMC to fit MSC model parameters to observed distances between tree-change and/or topology-change events in simulated ARGs.

Given one or more ARGs, each composing a sequence of genealogies G = (*𝒢*_1_, *𝒢*_2_, …, *𝒢*_*n*_) and their interval lengths in the genome X = (*x*_1_, *x*_2_, …, *x*_*n*_), a subset of genealogies can be extracted that represent the unique trees between tree-change events, G_*g*_ = (*𝒢*_1_, *𝒢*_*i*_, …), and waiting distances can be measured as the summed interval lengths between these trees, 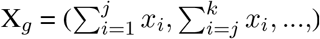. The same could be done for topology-change events. Given a parameterized species tree *𝒮* and recombination rate *r* we can then obtain a sequence of exponential rate parameters Λ_*g*_ = (*λ*_1_, *λ*_*i*_, …) corresponding to each genealogy in G_*g*_ (equations 10 or 14). The likelihood of each observed waiting distance in X_*g*_ can then be calculated from the exponential probability density function (equation 8) with rate parameters from Λ_*g*_. The maximum likelihood solution can be found by searching for the set of MSC model parameters – which affect Λ_*g*_ – that maximize the summed log-likelihood of the observed waiting distances.

We first implemented this approach for a simple two-population model with constant *N*_*e*_=200K and a population divergence at 500K generations. We simulated 100 independent ARGs, each 100Kb in length, using *r*=2e-9, and we sampled four haplotypes per population. This yielded 51,487 tree-change events, with mean length 194bp, and 25,446 topology-change events, with mean length 393bp. To examine the likelihood surface, we calculated the log-likelihood of joint parameters for *N*_*e*_ and *r*, while keeping a fixed divergence time parameter, over a grid search of 41 evenly spaced values from *r*=1e-9 to 3e-9 and *N*_*e*_=50K to 800K. The likelihood surface for tree-change distances exhibited a ridge containing the true parameter values (Fig. 5b), while the topology-change likelihood surface exhibited a distinct peak at the true values (Fig. 5c). The summed log-likelihoods from both tree and topology change distances provided the most informative likelihood surface (Fig. 5d). We also inferred a likelihood surface for a misspecified model, in which the genealogies were generated under a 2-population model, but the Λ_*g*_ rate parameters were calculated for a single population model with constant *N*_*e*_. This model misspecification introduced a significant bias in parameter estimation, shifting the likelihood surface towards lower *r* and higher *N*_*e*_ (Fig. 5e). This demonstrates that even for a simple two population model, ignoring population structure can significantly bias spatial genealogical inference methods.

**Figure 5.**
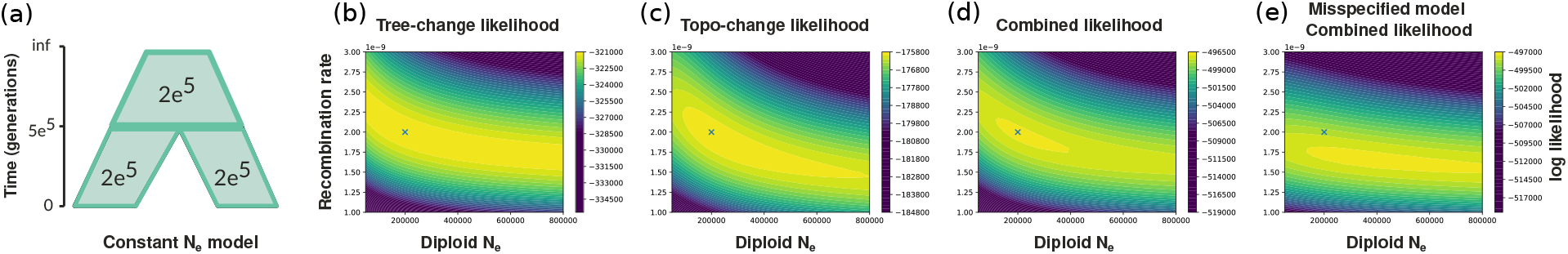
MS-SMC likelihood framework. (a) ARGs were simulated under a two-population species tree model with a constant *N*_*e*_=200K and *r*=2e-9. (b) A joint log-likelihood surface for *N*_*e*_ and *r* inferred from the distances between tree-change events; (c) topology-change events; or (d) both. The true parameters are marked by an X. (e) If the MSC model is misspecified as a single-population model but the data derive from a two-population model, likelihood inference is highly biased.

To examine parameter inference under more complex MSC models we implemented a Bayesian approach using a Metropolis-Hastings Markov chain Monte Carlo (MCMC) algorithm to compute the joint posterior probability distribution of multiple *N*_*e*_ parameters for a 2-population model with variable *N*_*e*_ of 200K, 300K, or 400K (Fig. 6a). The input ARGs were simulated under this more complex MSC model but otherwise used the same settings as in the example above. We set a uniform prior on all three *N*_*e*_ parameters between 1e^2^ and 1e^7^, and ran a single MCMC chain to sample 10K values, sampling every 5th iteration following a 1K iteration burnin. For simplicity, we fixed the population divergence time and recombination rate to their correct values to focus solely on *N*_*e*_ estimation. All parameters converged with ESS values exceeding 2000, and the 95% highest posterior density intervals (HPDI) for each posterior probability distribution included the true parameter (Fig. 6b). This suggests that multiple MSC model parameters can be accurately inferred from waiting distance information alone.

**Figure 6.**
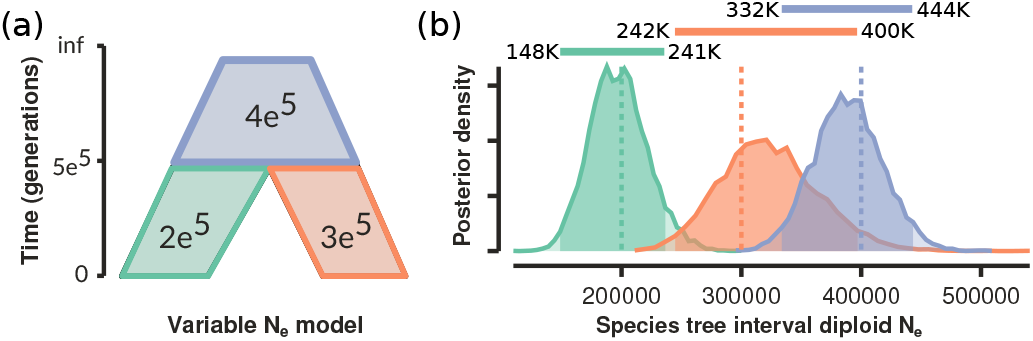
Joint inference of MSC model parameters using waiting distances calculated under the MS-SMC. (a) ARGs were simulated under a two-population MSC model with variable *N*_*e*_. (b) Posterior distributions of the three *N*_*e*_ parameters jointly inferred from tree- and topology-change waiting distances. The true model *N*_*e*_ values fall within the estimated 95% HPDI intervals.

## 4 Discussion

Genealogical relationships vary spatially across chromosomes, reflecting a history of recombination between genome segments inherited from different ancestors. Such variation can be modeled by the sequentially Markov coalescent, which provides a generative process upon which many statistical methods have been developed (McVean & Cardin, 2005; Spence *et al*., 2018). However, most applications of the SMC’ remain highly limited with regard to the scale over which they extract information from genomes – extending forward just one recombination event at a time. By contrast, the recent development of solutions for predicting tree- and topology-change waiting distances under the SMC’ (Deng *et al*., 2021) effectively adds two additional, longer-range sources of information for any position in a genome. In their model, these distances are a result of the probabilities of different phenomenological outcomes of the SMC’ process given a genealogy embedded in a single population coalescent model with constant *N*_*e*_. Here we extended this framework, deriving new solutions for the probabilities of tree-change and topology-change events for a genealogy embedded in any arbitrarily parameterized structured coalescent model. These solutions lay a groundwork for exploring how variation in species tree parameters affects neutral expectations of genealogical heterogeneity across chromosomes.

### 4.1 Applications of the MS-SMC

Our MS-SMC approach provides a predictive model for the relationship between a parameterized species tree and the length of a genomic interval over which a genealogy is expected to be observed. We demonstrated the accuracy of our implementation relative to stochastic simulations performed under both the full coalescent with recombination and the SMC’ approximation (Hudson, 1983; McVean & Cardin, 2005; Wiuf & Hein, 1999). Within our framework, MSC model parameters determine probabilities of coalescence in species tree intervals and prevent coalescence between lineages that are separated by species divergence events. The effects of MSC model parameters on spatial genealogical variation can be statistically modeled and visualized (Fig. 1; Fig. S2), and our examples show that the neutral rate of turnover in genealogical trees and topologies across a genome is highly dependent on species tree model parameters.

A complex relationship exists between a parameterized MSC model, the distribution of genealogies that can arise under that model, and the spatial distances over which those genealogies are expected to span. Previously, such patterns could only be examined through stochastic simulations. For example, McKenzie & Eaton (2020b) used exhaustive simulations to examine the effect of species tree parameters on genetic linkage by varying species tree length, size, and shape. While this approach is practical for estimating the *mean* linkage across a large set of sampled genealogies under a specific demographic model, it is impractical for estimating the persistence of *individual* genealogies. The analytical solutions presented here not only enable fast calculations but also the opportunity to develop likelihood-based methods for fitting models that link MSC parameters to ARGs.

Many extensions of the MS-SMC are possible, including applications to more complex demographic models and the development of specific hypothesis testing frameworks. For example, genealogies could instead be embedded in a species network following the multispecies network coalescent model (Wen *et al*., 2016). This would involve modeling the coalescent histories as a probabilistic process that could follow one of multiple possible paths through a network. Differences between the expected waiting distances in genomic regions derived from one embedding path history versus another may be highly informative about the network model in which genealogies are embedded. Therefore, incorporating linked genealogical information may prove valuable towards resolving intractable problems in species tree or network inference by moving beyond the limited information available from the frequencies of unlinked gene trees.

### 4.2 Phylogenetic and local ancestry inference

By linking species tree parameters to observable patterns in ARGs, the MS-SMC can be used to improve both gene tree and species tree inference methods. One practical application of waiting distance expectations calculated under the MS-SMC is to guide the selection of appropriate locus lengths for MSC analyses so as to avoid intra-locus recombination. This process yields data within a locus that derive from multiple variable genealogies, and so inferred gene trees represent concatenation artifacts. Topology-change waiting distances are often much shorter than sampled locus lengths for multispecies datasets (McKenzie & Eaton, 2020b), and this is especially likely when gene trees are inferred from large genomic sliding windows (e.g., Li *et al*., 2019). When repeated across many loci, concatenation artifacts can cause the distribution of gene trees, or of their summary statistics, to deviate from expectations under the MSC – a process termed concatalescence (Gatesy & Springer, 2013).

One way to improve gene tree inference for individual loci is to re-conceive the problem as one in which loci can have multiple gene tree histories and are thus better represented as ARGs. Applications of ARG inference have historically been applied at the scale of entire chromosomes, and usually to samples that are modeled as being in a single population (Kelleher *et al*., 2019; Rasmussen *et al*., 2014; Speidel *et al*., 2019). However, ARGs can exist for any stretch of the genome, including short aligned contigs, and recent methods can infer ARGs conditional on a structured demographic model that includes splits between more distantly related genomes (Hubisz & Siepel, 2020). Although our example likelihood-based implementation of the MS-SMC showed that MSC model parameters can be estimated from an observed ARG (Fig. 5, Fig. 6), we expect that many useful applications of the MS-SMC will likely come from the inverse application: inferring ARGs given a parameterized species tree model. Or, from joint inference of both MSC models and ARGs. For example, although ARGWeaver-D (Hubisz & Siepel, 2020) can currently propose an ARG conditional on a structured demographic model, which can generate genealogies with tree- and topology-change waiting distances as predicted under the SMC’, these distances do not contribute to the likelihood score of an ARG. We speculate that incorporating this information could improve both accuracy and convergence.

In contrast to current phylogenomic inference approaches which tend to either ignore genetic linkage, or to discard the vast majority of sequenced data in effort to avoid it, one could envision an alternative, spatially aware phylogenetic inference framework that more effectively utilizes linked genomic data. This would mark a major transition in phylogenetics, where recombination could be viewed as a source of information rather than a source of error. We see the MS-SMC as an important step in this direction. For example, a joint ARG and species tree inference approach might build upon the species tree inference method SNAPP (Bryant *et al*., 2012). SNAPP subsamples a single unlinked SNP from each locus and integrates over a distribution of genealogies that could have produced it, with the goal of bypassing the problem of gene tree inference. A theoretical ARG-based extension of this approach might analyze many linked SNPs and integrate over a distribution of ARGs that could have produced each linked pattern, with the goal of bypassing ARG inference. By reducing the complexity of ARG inference to a problem of windows between topology-change events, as opposed to any two recombination events, the MS-SMC provides a more efficient framework that could potentially make methods linking ARGs and species tree inference possible.

Finally, a common goal of many local ancestry inference methods is to test evolutionary hypotheses involving selection or introgression (Martin & Belleghem, 2017). For example, sliding window analyses have been used to identify putatively introgressed loci in *Heliconius* butterflies (Zhang *et al*., 2016) and felids (Li *et al*., 2019). Such conclusions are often based on the genomic interval lengths over which certain topological gene tree patterns are observed. We suggest that such conclusions should be examined critically when made without reference to a null-model-based expectation. Our results show that neutral expectations for waiting distances between genealogy changes can exhibit high variance, and that this variance will be spatially auto-correlated, such that a topology or clade may exist over long stretches of the genome simply by chance.

## 4.3 Acknowledgements

This work was supported by the National Science Foundation (NSF DEB-2046813 awarded to D.A.R.E. and NSF Graduate Research Fellowship DGE 16-44869 awarded to P.F.M.). Thanks to Yun Deng for discussion on waiting distance methods, and to members of the Eaton Lab for valuable feedback.

## 5 Supplementary Information

### 5.1 Supplementary Figures

**Figure S1.**
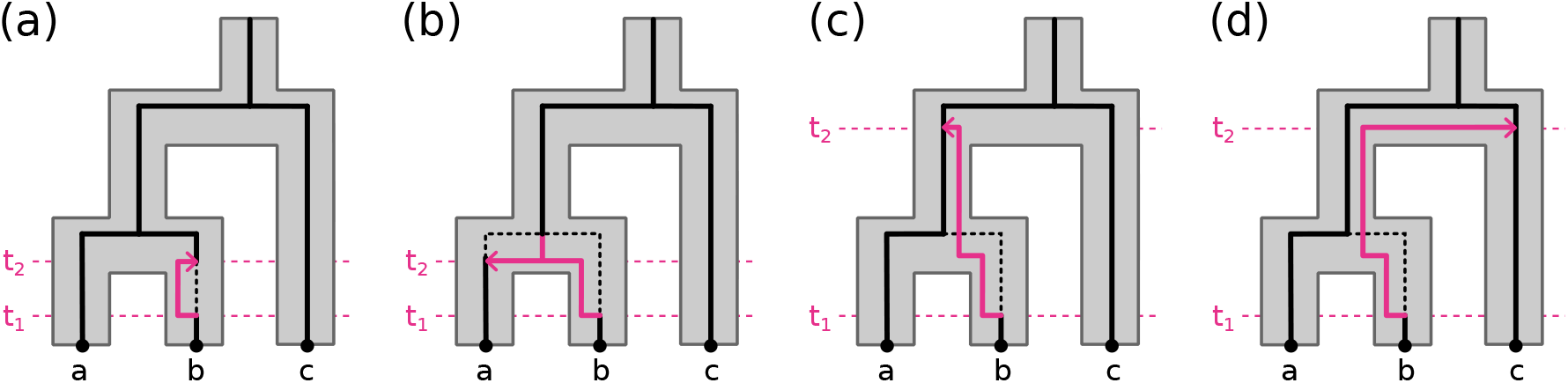
Four categories of outcomes from a recombination event occurring on a genealogy at time t_1_, dictated by random subtree re-coalescence with a remaining lineage under the SMC’ process at time t_2_. (a) The detached subtree re-coalesces with the original lineage from which it was detached, leading to no change between the starting genealogy and subsequent genealogy. (b) The detached subtree re-coalesces with its sibling lineage prior to their previous coalescence, leading to a shortening of their coalescence time. (c) The detached subtree re-coalesces with its parent lineage, leading to a lengthening of the coalescent time between the detached subtree lineage and its sibling lineage. (d) The detached subtree re-coalesces with a lineage other than itself, its sibling, or its parent lineage, leading to a topology-change.

**Figure S2.**
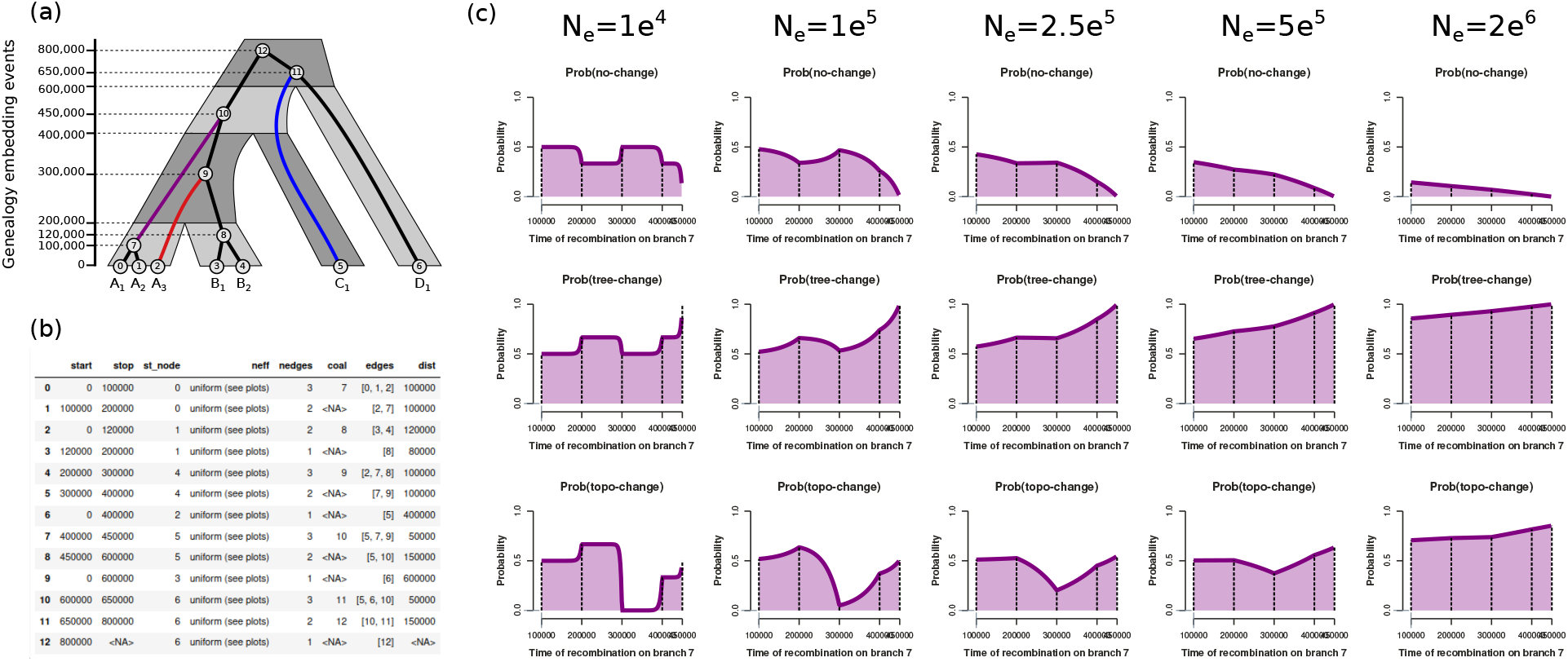
Probabilities of different recombination event outcomes for a selected genealogy edge as a function of the time at which recombination occurs and of the constant effective population size. (a) An MSC model with edge lengths in units of generations and an example genealogy embedded. (b) An genealogy embedding table for the example MSC model and genealogy. (c) Probabilities of different recombination event outcomes across genealogy edge 7. When *N*_*e*_ is low, probabilities are nearly constant with respect to time within each interval since re-coalescence in later intervals is unlikely. When *N*_*e*_ is high, probabilities change nearly monotonically across the length of an edge since population structure does little to constrain the time of re-coalescence.

### 5.2 Investigating bias in MS-SMC predictions

The MS-SMC harbors two potential sources of bias, the first stemming from assumptions of the SMC’ approximation and the second from the approximate nature of waiting distance estimation for topology-change events. We examined both of these sources of error through comparison to stochastic simulations and found that their effects are generally negligible and act in opposing directions such that their combined effect is further reduced.

The SMC’ approximation to the full coalescent with recombination is expected to deviate increasingly far from the full model as the number of recombination events that the SMC’ does not model (i.e., those that are among ancestors who do not contribute genetic material to sampled descendants) increases. By not modeling this subset of recombination events, the SMC’ will tend to exhibit greater waiting distances between tree or topology change events than the full model. To investigate this source of error in our waiting distance predictions, we repeated the simulations from our validation scenario using the *msprime* setting ancestry=“smc_prime” to simulate tree sequences while excluding recombination events that would not occur in the SMC’ model. We have already seen from our validations that the error in our predictions is quite low when data are simulated under the full coalescent with recombination (Fig. S3). As expected, we observe even less error between our predictions and coalescent simulations when data are simulated under the SMC’ model. Whereas the error rate is generally below 5% when data are simulated under the full coalescent with recombination model (only exceeding this at the lowest *N*_*e*_ values examined) mean error rates do not exceed 5% in any models for data simulated under the SMC’ assumptions. This shows that the SMC’ assumption does contribute a relatively small error to our waiting distance predictions, especially at low *N*_*e*_ values, where it can lead to over-estimated waiting distances.

We also investigated whether inhomogeneity in the probability of topology changes during the waiting distance between topology-change events causes bias. For this, we employed a similar approach to Deng *et al*. (2021), who examined the fold-difference in parameters affecting waiting distance estimations between a starting tree and a subsequent genealogy that experienced a tree-change but not a topology-change. Because MSC model parameters affect both the length of genealogies and the probability of topology-change, we also examined variation in each of these parameters at different constant *N*_*e*_ values of 50K, 100K, or 500K. For each setting we examined one topology-change event from 1K tree sequences.

In a single population model with constant *N*_*e*_, Deng *et al*. (2021) previously showed that the bias in topology-change waiting distances is negligible because of an inverse relationship between the length of a genealogy and the probability of a topology change. Our analysis confirms this result, showing that variance in the product of these two parameters, which equates to the waiting distance rate parameter (equations 7, 8, 10), is very low (Fig. S4a) and becomes smaller as more gene copies are sampled (Fig. S4b-c). In the 2-population and 8-population MSC-type models we find the same result. Here, constraints imposed by population structure lead to less variance in both the length of genealogies and probabilities of topology-change (Fig. S4d-i). When *N*_*e*_ is low, there is very little variation between subsequent genealogies, and thus the waiting distance expectation exhibits little heterogeneity. When *N*_*e*_ is high, there is little population structure, and so the waiting distance expectation remains relatively constant for the same reason as in a single population model. Overall, MSC models do not appear to exhibit a greater bias from this source than single population models.

**Figure S3.**
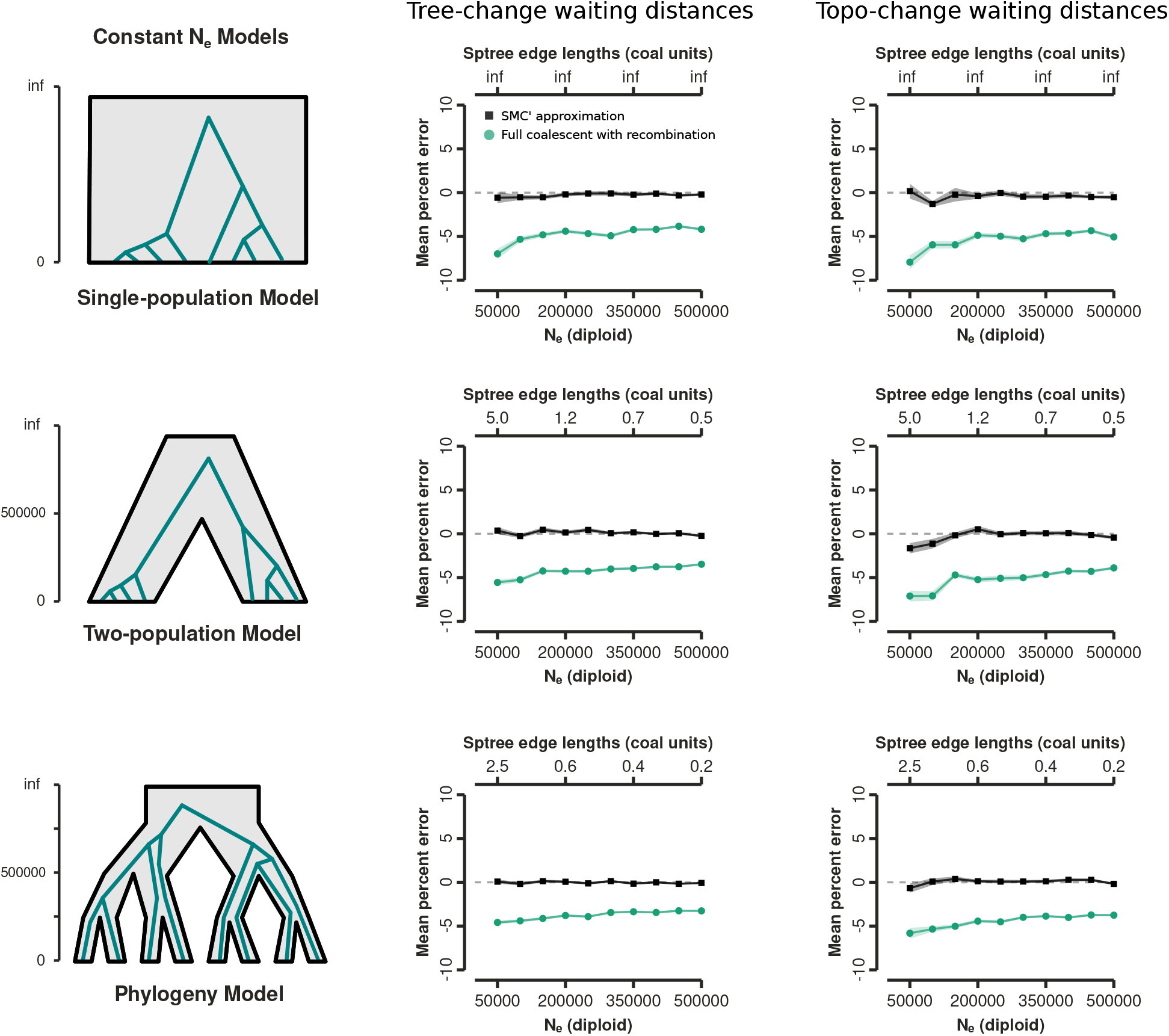
Error in MS-SMC waiting distance expectations caused by the SMC’ approximation. Error was measured as the mean percent difference between expected waiting distances calculated under the MS-SMC and observed waiting distances in stochastic coalescent simulations. Simulations were performed under either the SMC’ approximation (black) or the full coalescent with recombination (green). The MS-SMC tends to under-estimate waiting distances compared to the full coalescent with recombination but shows only a slight bias at very low *N*_*e*_ values compared to simulations under the SMC’.

**Figure S4.**
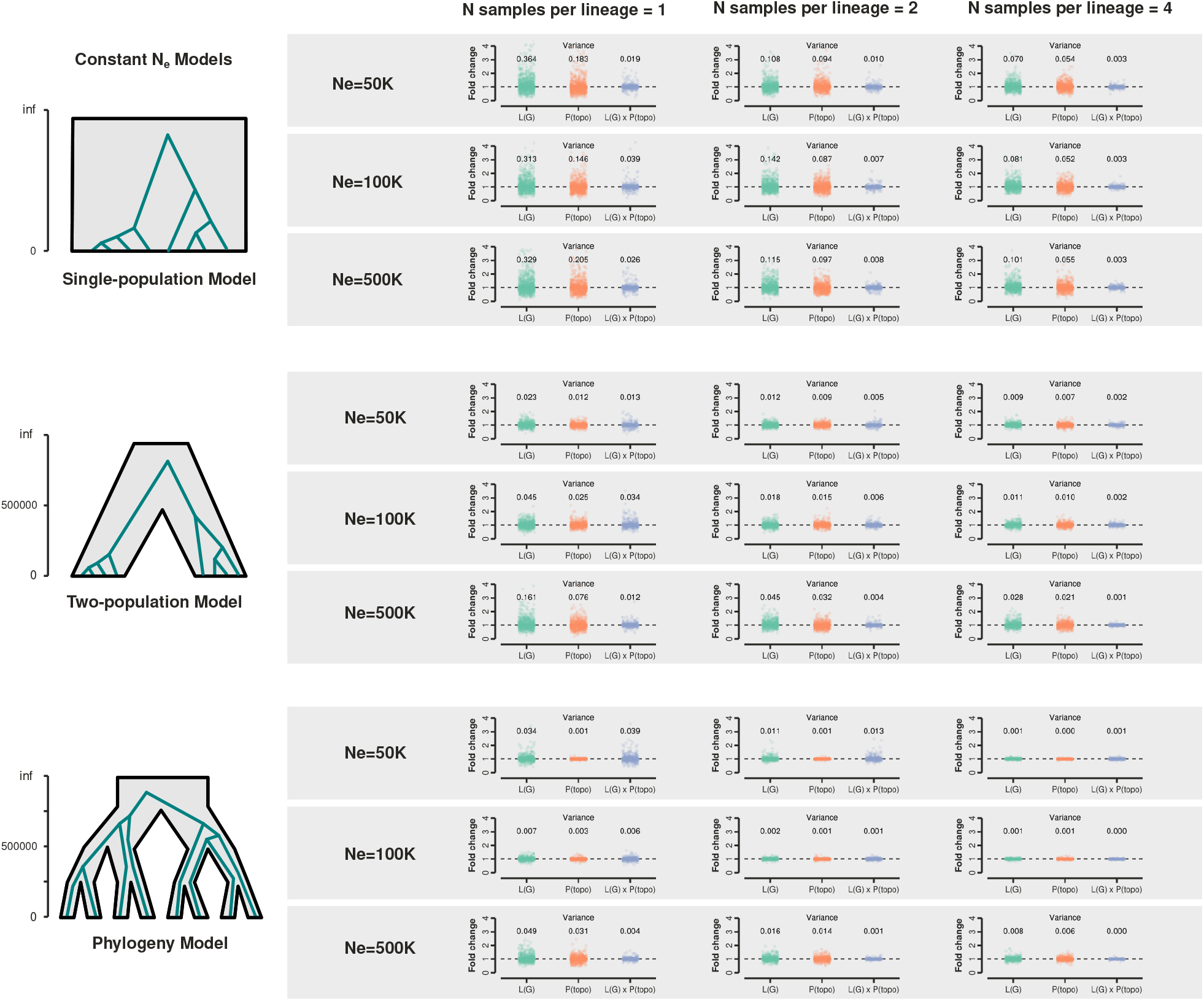
Variance in the fold-change for components affecting the expected waiting distance to a topology-change event between the starting tree and a subsequent tree which has experienced a tree-change event, changing the coalescent times but not the topology. The sum of genealogical edge lengths (L(G)), the P(topology-change | S,G), and the product of these two terms are shown for three different demographic models and with different constant *N*_*e*_ values, and/or numbers of samples per lineage. When the fold-change in the product exhibits low variance around 1 the MS-SMC approximation for the expected waiting distance until a topology-change is expected to be more accurate. Larger effective population sizes and numbers of samples per lineage yield lower variance in the product.

## 6 Appendix: Derivations

### 6.1 Notation

Information from the genealogy embedding table (described in the following paragraph) can be used in equations that calculate the probabilities of no-change, tree-change, and topology-change events under the MS-SMC. These equations, described throughout rest of the Appendix, use the terms defined in Table S1. A parameterized species tree, *𝒮*, is a multispecies coalescent model in which a set of isolated populations are related by a bifurcating tree topology. Divergence times between lineages are in units of generations, and each edge (species tree interval) can be associated with a different constant diploid effective population size (*N*_*e*_). A genealogy, *𝒢*, represents the genealogical relationships – composing a topology and coalescent times in units of generations – for a set of sampled gene copies at some position in their genomes. A genealogy can be embedded in a species tree if the coalescent times between sampled gene copies from different populations are not younger than a population divergence event separating them.

Given a genealogy embedded in a species tree, a series of discrete time intervals can be defined that are delimited by events that change the rate of coalescence. We refer to this set of discrete time intervals and their associated properties as a genealogy embedding table (e.g., Table 1). In the waiting distance solutions for a single population with constant *N*_*e*_ by Deng *et al*. (2021), this table is delimited only by coalescent events, and the intervals are non-overlapping. Because *N*_*e*_ is constant in their framework, only *k* differs between intervals. Therefore, changes in *k* alone determine differences in rates of coalescence, with *k* decreasing monotonically in subsequent intervals from the tips towards the root. Our approach is similar, but adds additional complexity (Fig. 1). In the multispecies framework, genealogy embedding intervals are specific to each species tree branch, with each one corresponding to a time interval with a constant *k* and *N*_*e*_ in a specific species tree branch. Breakpoints between intervals arise where divergences occur in the species tree (increasing *k* and potentially changing *N*_*e*_) and where coalescent events occur in the genealogy (reducing *k*). Genealogy embedding intervals corresponding to different species tree branches can overlap in time.

**Table S1.** Summary of variables used in waiting distance equations.

| Variable | Description |
| --- | --- |
| $\mathcal{S}$ | An MSC model with topology, divergence times and effective population sizes. |
| $\mathcal{G}$ | A genealogy that can be embedded in $\mathcal{S}$. |
| $L(\mathcal{G})$ | Sum of edge lengths of genealogy $\mathcal{G}$. |
| $b$ | A focal branch in $\mathcal{G}$. |
| $i$ | Interval in the genealogy embedding table in which recombination occurs. |
| $\mathcal{I}_b$ | Ordered set of intervals on branch $b$. |
| $\mathcal{I}_c$ | Ordered set of intervals on branch $c$, the parent of branch $b$. |
| $\mathcal{I}_{bc}$ | Ordered union of sets $\mathcal{I}_b$ and $\mathcal{I}_c$. |
| $\mathcal{J}_b(i)$ | Ordered set of intervals above $i$ on branch $b$. |
| $\mathcal{Q}_b(i, j)$ | Ordered set of intervals above $i$ and below $j$ on branch $b$. |
| $\mathcal{K}(b, t)$ | Number of edges of $\mathcal{G}$ in the interval containing branch $b$ at time $t$. |
| $k_x$ | Number of edges of $\mathcal{G}$ in interval $x$; piece-wise constant of $A(b, t)$. |
| $\mathcal{N}(b, t)$ | Diploid effective population size in the interval containing branch $b$ at time $t$. |
| $n_x$ | Diploid effective population size in interval $x$; piece-wise constant of $N(b, t)$. |
| $t_r$ | Time of a recombination event, in generations. |
| $\sigma_x$ | The lower boundary of interval $x$, in generations. |
| $\mu_x$ | The upper boundary of interval $x$, in generations. |
| $d_x$ | The length of interval $x$, in generations. |
| $t_b^l$ | The lower boundary of branch $b$, in generations. |
| $t_b^u$ | The upper boundary of branch $b$, in generations. |
| $t_b^m$ | The time at which a focal branch $b$ is able to coalesce with its sibling branch. |
| $\mathcal{M}_b$ | Ordered set of intervals above $t_b^m$ on branch $b$. |
| $\mathcal{L}_b$ | Ordered set of intervals below $t_b^m$ on branch $b$. |

Each branch on *𝒢* will span one or more genealogy embedding intervals. The ordered set of intervals on a specific branch, *b*, is defined as *ℐ*_*b*_. The lower and upper time bounding each interval is *σ*_*x*_ and *μ*_*x*_, respectively, where *x* is the index of the interval in the genealogy embedding table. The lower and upper bounds of each branch are defined as 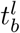 and 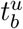, respectively.

### 6.2 Extending SMC’ waiting distance solutions

The probabilities of different recombination event types under the MS-SMC are calculated from the probability that recombination occurs on a specific branch and the probabilities that the resulting detached subtree subsequently re-coalesces with any other available branch above that time. The opportunity for recombination to occur on a branch is scaled by its length in generations 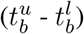. Similarly, the probability of re-coalescence on a branch is scaled by its length and the coalescence rate. The latter can vary over the length of a branch as it spans different intervals, and is a function of the effective population size in the species tree interval that includes branch *b* at a specified time, *τ*, defined as *𝒩*(*b, τ*), and the number of other genealogy branches in the interval that includes branch *b* at time *τ*, defined as *𝒦*(*b, τ*). Finally, the probability that coalescence occurs over an interval of length (*t*) can be calculated from an exponential probability density *f* (*t*; *λ*), where the rate parameter is 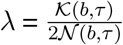, similar to equations 1-2.

### 6.3 Probability of no-change

#### 6.3.1 Given a branch and time of recombination

The probability that a tree is unchanged by a recombination event – meaning that no coalescent times are changed – is the probability that the detached subtree re-coalesces with the same branch it detached from. Thus, we can integrate from the time of recombination (*t*_*r*_) to the top of the branch 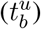 over the probability of sampling the same branch times the exponential probability density of re-coalescing at any time on that branch above the time of recombination (*τ* − *t*_*r*_). We take this integral with respect to *τ*, where *𝒦*(*b, τ*) and *𝒩*(*b, τ*) can vary across the length of the branch if it spans different intervals.

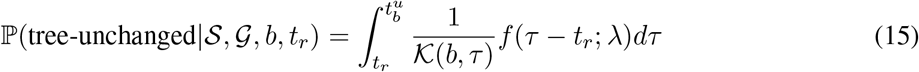

The exponential probability density function can be expanded, as in equation 2, where *λ* is *𝒦*(*b, τ*) over 2*𝒩*(*b, τ*):

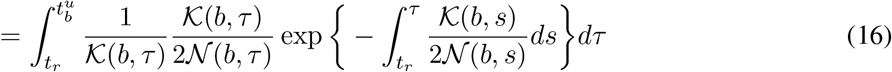

This simplifies to the following equation, which describes the probability that the subtree re-coalesces at any time (*dτ*) on *b* above *t*_*r*_, and that it does not re-coalesce at any intervening time (*ds*) between *t*_*r*_ and *τ*.

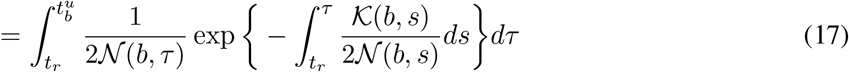

Because the rate of re-coalescence is constant within each interval, we next split this equation into statements over each discrete interval that the detached subtree could possibly re-coalesce with on branch *b*. (Recall, because we are currently computing the probability of a no-change event we only need to concern ourselves with re-coalescence on branch *b*.) Here, the interval in which recombination occurred on branch *b* is labeled as *i*.

In the equation below, the first integral describes the probability statement over only part of interval *i*, from *t*_*r*_ to *u*_*i*_, rather than over its entire length, since re-coalescence can only occur above the time at which recombination occurred. By contrast, the latter parts of this equation are performed over the entire lengths of each remaining interval above *i*, from the bottom (*σ*_*j*_) to the top (*μ*_*j*_) of the interval. The ordered set of all intervals above *i* in *ℐ*_*b*_ is defined as *𝒥*_*b*_(*i*) = *{j* ∈ *ℐ*_*b*_ | *j > i}*.

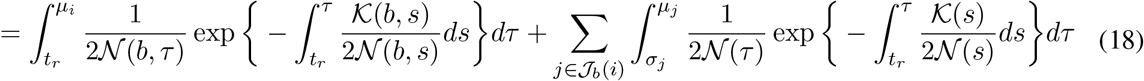

We can now solve this equation and substitute constant values for *𝒦*(*b, τ*) and *𝒩* (*b, τ*) in each interval. Because the first term is computed over only part of the first interval we first solve this term separately, and then show the result for the later terms. The first term concerns the probability of re-coalescing in the same interval *i* in which recombination occurred. The center part of this equation will appear again later, and so we define it as the function *f* (*i, i*).

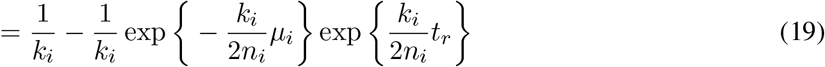

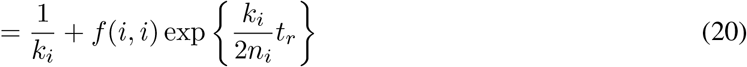

We similarly define the function *f* (*i, j*) for the later terms in this equation, which refer to the probability of re-coalescing in a later interval, *j*, than the one in which recombination occurred, *i*. This function requires also summing over any intervening intervals, *q*. For this, we define the function *𝒬*_*b*_(*i, j*) = *{q* ∈ *ℐ*_*b*_ | *j > q > i}* to return the ordered set of intervals on branch *b* between *i* and *j*:

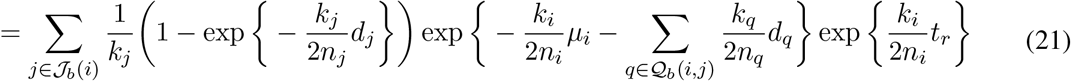

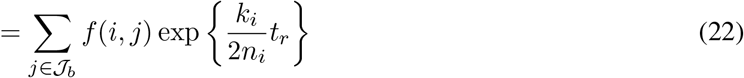

The *f* (*i, i*) and *f* (*i, j*) function above is represented in the main text as equation 4. Adding the two terms that include these functions together we get equation 3 from the main text for the probability of a nochange event given the timing and branch on which recombination occurs:

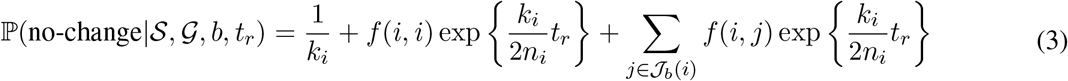

Note that in equation 3 above, while we have split the terms for clarity, the *f* (*i, i*) term could be lumped into the summation. The summation would then be indexed using *j* ∈ *ℐ*_*b*_, with the understanding that *f* (*i, j*) = 0 when *i > j* – that is, when summing over intervals that fall below the time of recombination.

#### 6.3.2 Across a full branch

Having solved for the probability of the genealogy being unchanged given the time *t*_*r*_ of the recombination event, our next step is to integrate this equation across the entire branch with respect to *t*_*r*_:

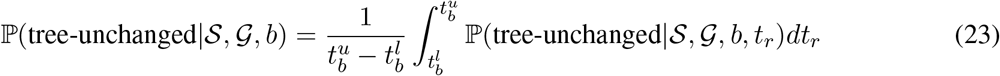

Plugging in the piece-wise constant solutions from equation 3 we get the following solution.

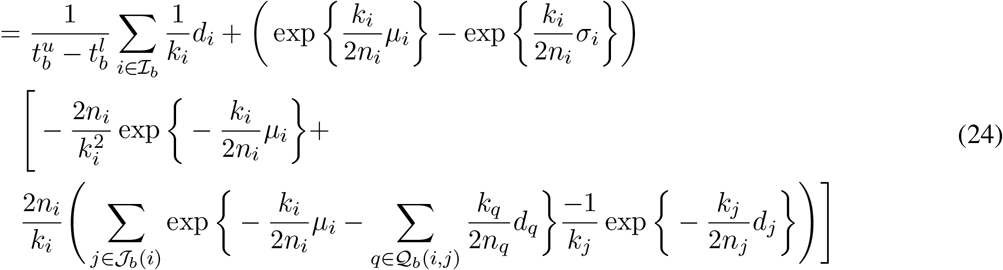

Finally, this can be simplified to the following solution by expressing the piecewise constant re-coalescence rates using the function *f* (*i, j*), as shown in equation 5 from the main text.

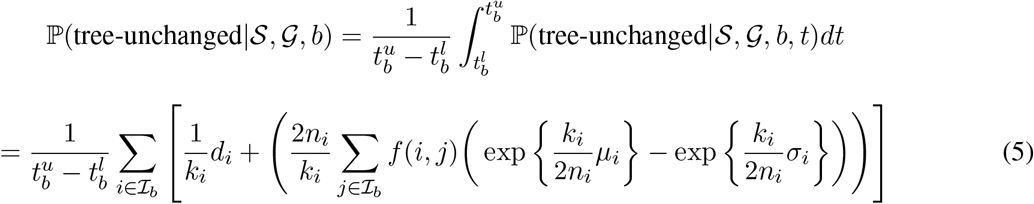

#### 6.3.3 Across the whole tree

At last, we can calculate the probability that, given a recombination event, the genealogy is unchanged. We do this by weighting each branch by its proportion of the total tree length and summing across the unchanging probabilities for all branches. This is equation 6 from the main text:

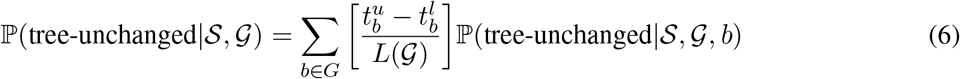

**Figure S5.**
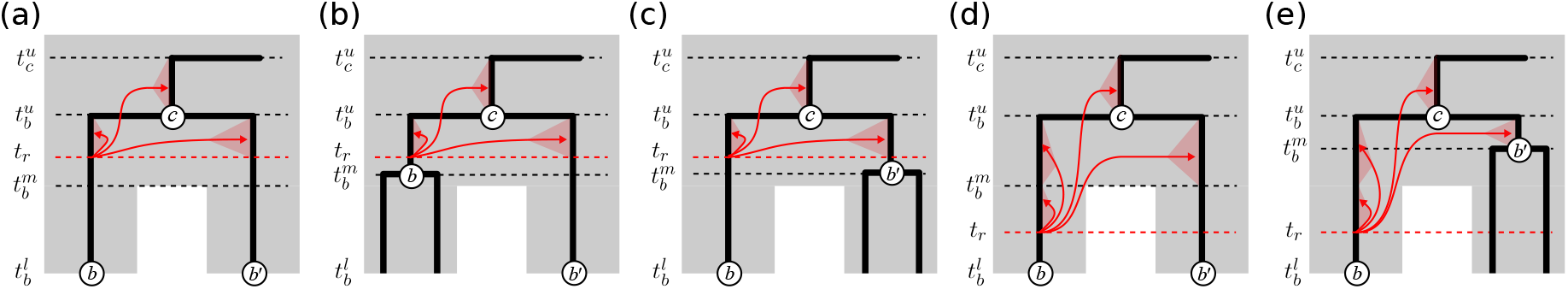
Calculating the probability that recombination on genealogy branch *b* leads to a topology change involves summing over the probabilities that the detached lineage does not re-coalesce with either itself, its sibling, or its parent (*b, b*^*′*^ or *c*, respectively). The possibility of a tree-change outcome (e.g., shortened coalescent time) is restricted until the lowest shared interval between *b* and *b*^*′*^, designated at time *t*_*m*_. Opportunities for such events could be constrained by species divergences – as in (a) and (d) – or by the timing of prior coalescence events generating each branch – as in (b), (c), and (e). The possibility that recombination (*t*_*r*_) occurs prior to *t*_*m*_ leads to the two ordered sets of intervals used in equations 11 and 12.

We can then also derive the probability of a tree-change event as 1 − ℙ(tree-unchanged| *𝒮, 𝒢*). Following the approach described in equations 7-10, we can then calculate an exponential probability distribution for waiting distances between no-change or tree-change events using an exponential rate parameter that is scaled by the probabilities of either recombination event type.

### 6.4 Probability of topology change

Next, we derive the probability of a topology-unchanged event, from which we can get the associated probability of a topology-change event. As in the first section, we first derive a solution given an individual branch and time of recombination. We then extend this solution to an entire branch, and we finally sum across branches to get a probability for the entire genealogy. To isolate events that do not cause a topology change, we must find the union of events that cause a no-change event in addition to two types of possible tree-change events which affect only branch lengths but not the topology. These two types of events correspond to a re-coalescence with the sibling to branch *b*, termed *b*^*′*^, or with its parent, *c* (Fig. 3d). Because branch *c* is always ancestral to *b*, a recombination event on *b* can potentially re-coalesce anywhere on *c*; however, this is not the case for *b*^*′*^, which may only exist or be available for re-coalescence over part of the length of *b*. It is therefore important to define the lowest time point at which re-coalescence with *b*^*′*^ is possible, termed 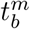.

In the single population model of Deng *et al*. (2021), 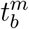 occurs at the maximum value of 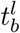 and 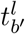, representing the lower bounds of *b* and *b*^*′*^, respectively. In our MSC-based model we must also incorporate potential constraints imposed by population barriers, and so 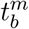 occurs at the maximum of 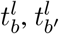, and any population divergence events that separate *b* and *b*^*′*^ (Fig. S5a-c).

#### 6.4.1 Given a branch and time of recombination

Using the definition for 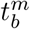 from above, we can describe the probability of a topology-unchanged event given the branch and time of recombination as a two-part solution. These two parts correspond to scenarios in which the time of recombination, *t*_*r*_, occurs either above (Fig. S5a-c) or below (Fig. S5d-e) 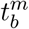. When 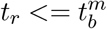, there are only two distinct intervals over which re-coalescence can occur: from *t*_*r*_ to 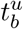 on branches *b* or *b*^*′*^, and from 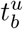 to 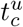 on branch *c* (Fig. S5a-c). By contrast, when 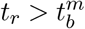 there are three distinct intervals for re-coalescence: from *t*_*r*_ to 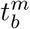 on *b*, 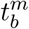 to 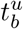 on *b*^*′*^, and 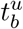 to 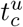 on branch *c* (Fig. S5d- e). Thus, in the first scenario the opportunity for re-coalescence is the same for branches *b* and *b*^*′*^, whereas in the latter scenario it is different.

Retaining the correct order of intervals is important in these calculations, particularly for *f* (*i, j*), which involves summing over not only the information in the *i* and *j* intervals, but also all of the intervals that lie between them. To iterate over ordered intervals on each branch we define additional indexing variables. Just as *ℐ*_*b*_ defines the ordered set of intervals on branch *b, ℐ*_*c*_ is the ordered set of intervals on branch *c*, and *ℐ*_*bc*_ is the ordered union of these sets. In addition, we define *ℳ*_*b*_ as the ordered intervals on branch *b* above 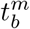, and *ℒ*_*b*_ as the ordered intervals on branch *b* below 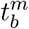. We can now derive a probability for the two distinct scenarios:

##### First case

Given 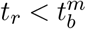, we integrate over the three distinct intervals where re-coalescence can occur. The first is unique to branch *b*, the second integral is multiplied by two since from 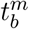 to 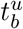 re-coalescence can occur with *b* or *b*^*′*^, and the final integral is over the length of branch *c*. By substituting the piece-wise constant solutions for each interval into this equation it can simplifed to the final form below, also shown in equation 11 of the main text:

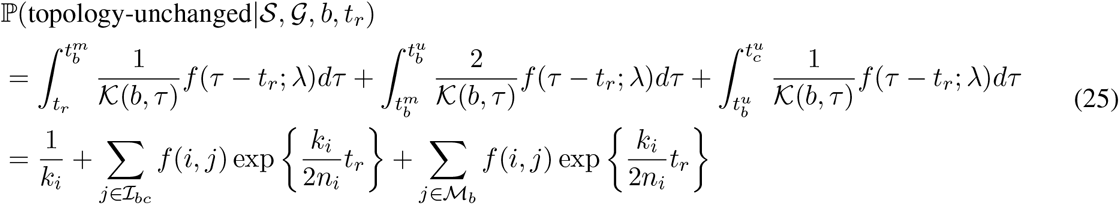

##### Second case

Given 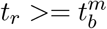, we only need to integrate over two distinct intervals, thus we simply drop the first term from the equation above. The final form of this equation is also shown in equation 11 of the main text:

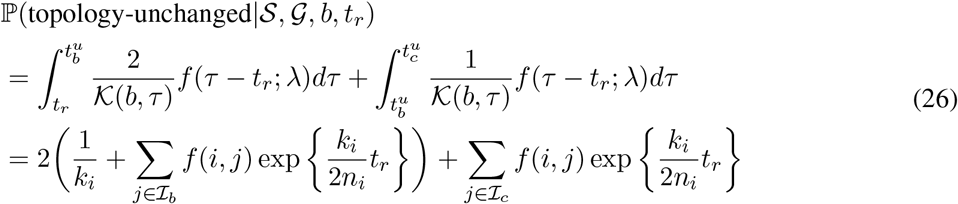

#### 6.4.2 Across a full branch

Now we derive the overall probability that a recombination event falling on a specific branch will change the topology by integrating across the range of possible values for *t*_*r*_. Following the approach above, we split this problem into two parts, above and below 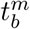, and we sum the two cases.

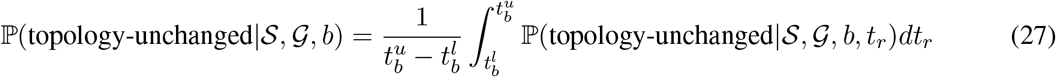

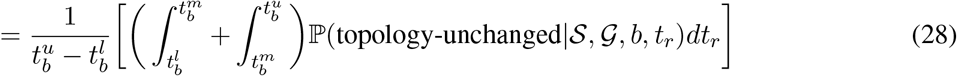

##### First case

We can simply sum over each entire interval below 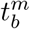 where *t*_*r*_ could occur, and substitute piece-wise constant solutions for the probabilities that a detached subtree will re-coalesce over the subset of targeted intervals above this that do not cause a topology-change.

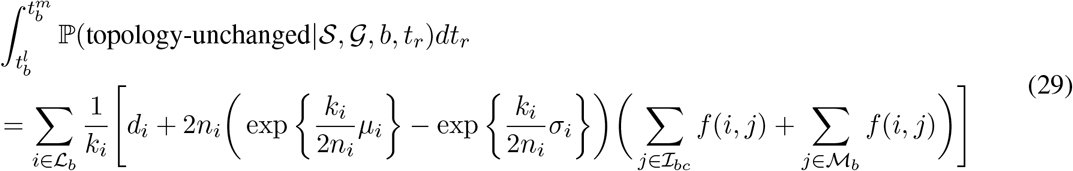

##### Second case

Similarly, we can sum over each entire interval above 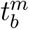 (up to 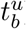) and substitute piece- wise constant solutions for the same selected subset of intervals:

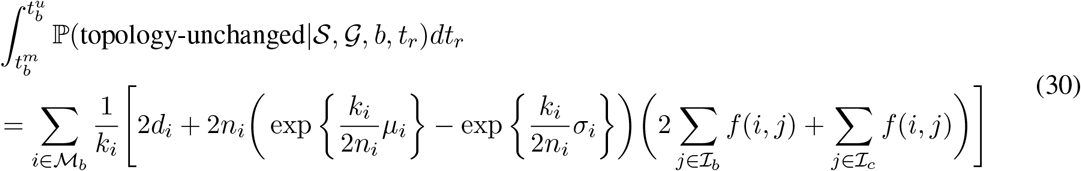

##### Result

If we express the inner summed terms from the equations above, composed of piece-wise constant values from their intervals, as 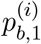 and 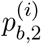, respectively, then the final solution can be expressed more concisely. This is shown in the main text as equation 12.

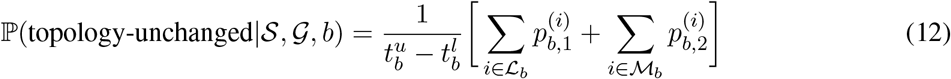

#### 6.4.3 Across the whole tree

Finally, we sum across all branches, each weighted by their relative length, to find the probability of a recombination event changing the topology of the tree. This appears as equation 13 in the main text.

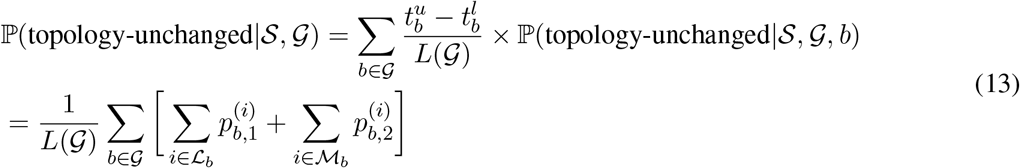

#### 6.4.4 Examples

Examples showing how to compute the probablity of a no-change (tree-unchanged) or topology-change event are shown with didactic step-by-step instructions in Fig. S6 and Fig. S7, respectively.

**Figure S6.**
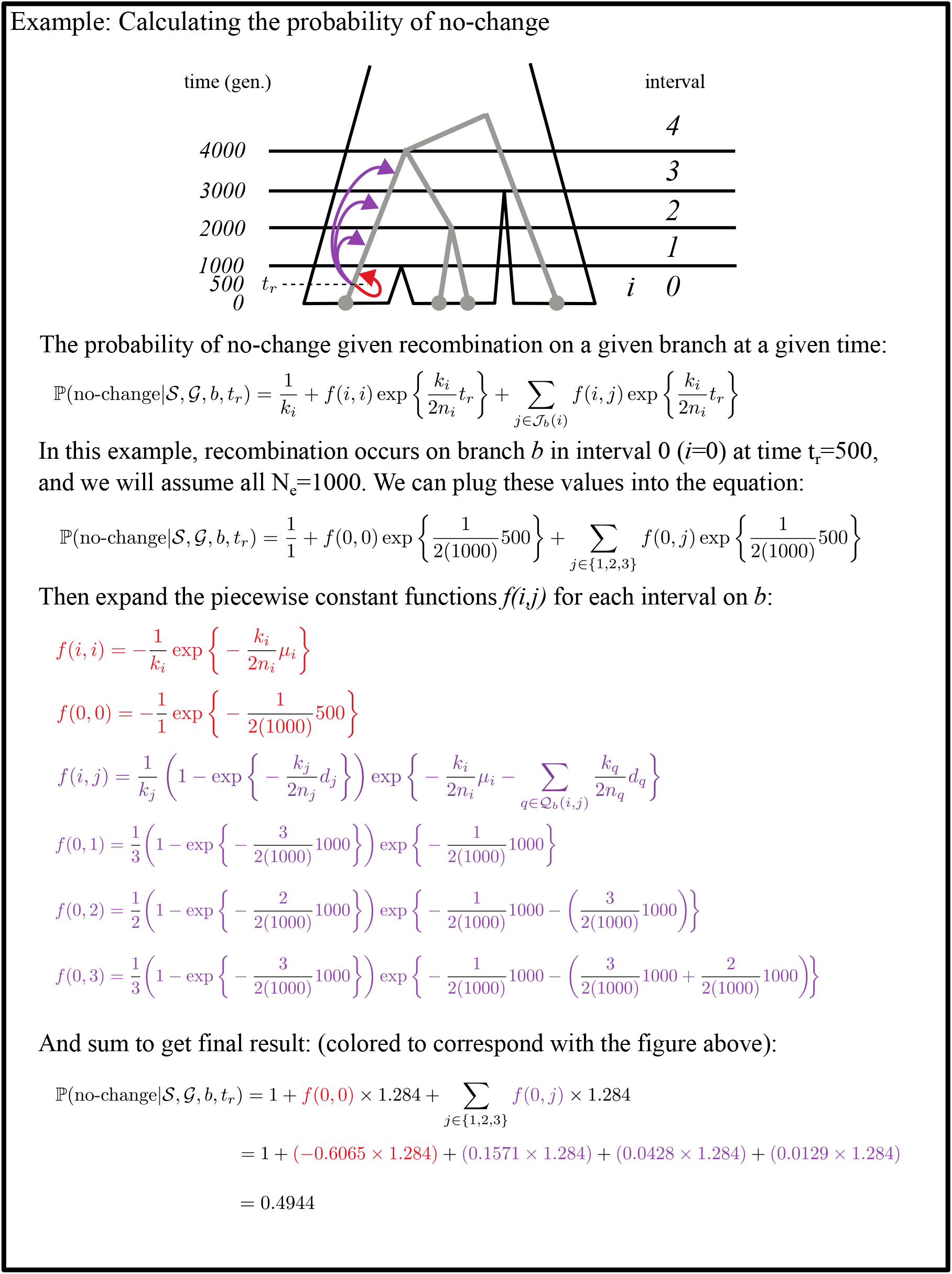
A step-by-step calculation of the probability of a tree-unchanged event under the MS-SMC given a species tree and genealogy.

**Figure S7.**
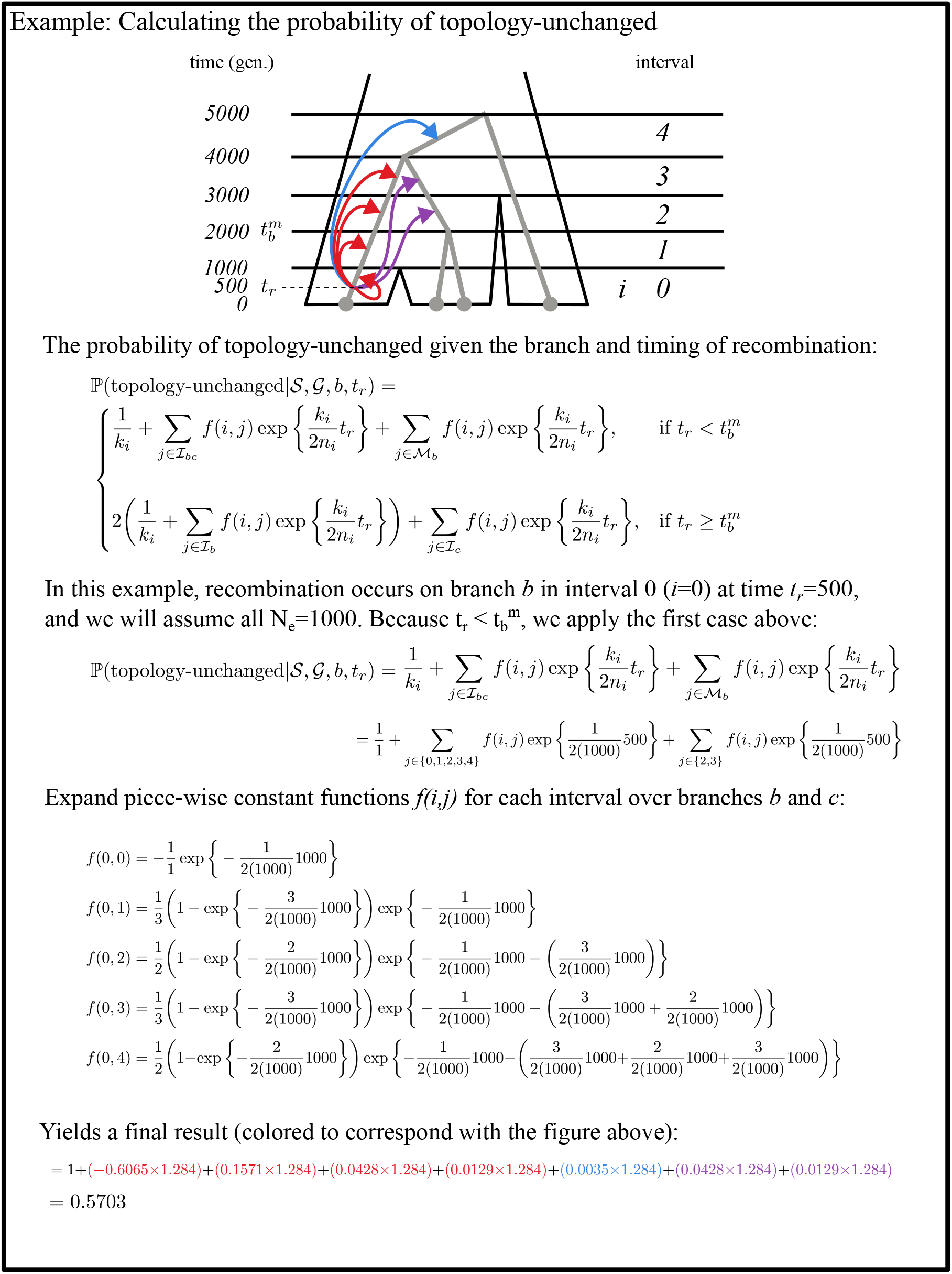
A step-by-step calculation of the probability of a topology-unchanged event under the MS-SMC given a species tree and genealogy.

